# FliH and FliI help FlhA bring strict order to flagellar protein export in *Salmonella*

**DOI:** 10.1101/2023.11.25.568686

**Authors:** Miki Kinoshita, Tohru Minamino, Takayuki Uchihashi, Keiichi Namba

**Affiliations:** Graduate School of Frontier Biosciences, Osaka University, 1-3 Yamadaoka, Suita, Osaka 565-0871, Japan; Department of Physics, Nagoya University, Chikusa-ku, Nagoya 464-8602, Japan; JEOL YOKOGUSHI Research Alliance Laboratories, Osaka University, 1-3 Yamadaoka, Suita, Osaka 565-0871, Japan

## Abstract

The flagellar type III secretion system (fT3SS) switches substrate specificity from rod-hook-type to filament-type upon hook completion, terminating hook assembly and initiating filament assembly. The FlhA ring is directly involved in substrate recognition, allowing the fT3SS to coordinate flagellar protein export with assembly, but the mechanism remains a mystery. Here, we report that the highly conserved GYXLI motif of FlhA is important for ordered protein export by the fT3SS. The fT3SS with the *flhA(Y369A/R370A/L371A/I372A)* (AAAA) or *flhA(Y369G/R370G/L371G/I372G)* (GGGG) mutation did not switch the substrate specificity at an appropriate timing of hook assembly. The A372V/T and G372V substitutions recovered the export switching function of the AAAA and GGGG mutants, respectively, in the presence but not in the absence of FliH and FliI, components of the flagellar ATPase complex. Interestingly, a filament-type substrate, FlgL, was secreted via the fT3SS with the AAAA or GGGG mutation during hook assembly in the absence of FliH and FliI but not in their presence. These observations suggest that FlhA requires the flagellar ATPase complex not only to efficiently remodel its ring structure responsible for the substrate specificity switching of the fT3SS but also to correct substrate recognition errors that occur during flagellar assembly.

## Introduction

*Salmonella enterica* serovar Typhimurium (hereafter referred to as *Salmonella*) can swim in liquids and move on solid surfaces by rotating flagella. The *Salmonella* flagellum consists of the basal body that acts as a bi-directional rotary motor fueled by proton motive force across the cytoplasmic membrane, the filament that functions as a helical propeller to produce propulsive force, and the hook that works as a universal joint to smoothly transmit torque produced by the motor to the helical propeller. Flagellar assembly begins with the basal body, followed by the hook and finally the filament (Fig. 1)^1,2^. To construct the flagellum on the cell surface, the flagellar type III secretion system is located at the base of the flagellum and transports flagellar structural subunits from the cytoplasm to the distal end of the growing flagellar structure. The fT3SS consists of a transmembrane export gate complex made of FlhA, FlhB, FliP, FliQ, and FliR and a cytoplasmic ATPase complex consisting of FliH, FliI, and FliJ (Fig. 1) ^3,4^. The export gate complex uses both H^+^ and Na^+^ as the coupling ion and acts as a cation/protein antiporter that couples inward-directed cation flow with outward-directed protein translocation^5–11^. The cytoplasmic ATPase complex acts not only as an ATP-driven activator of the export gate complex but also as a dynamic carrier that delivers export substrates and chaperone-substrate complexes from the cytoplasm to the export gate complex^12,13^.

**Fig. 1.**
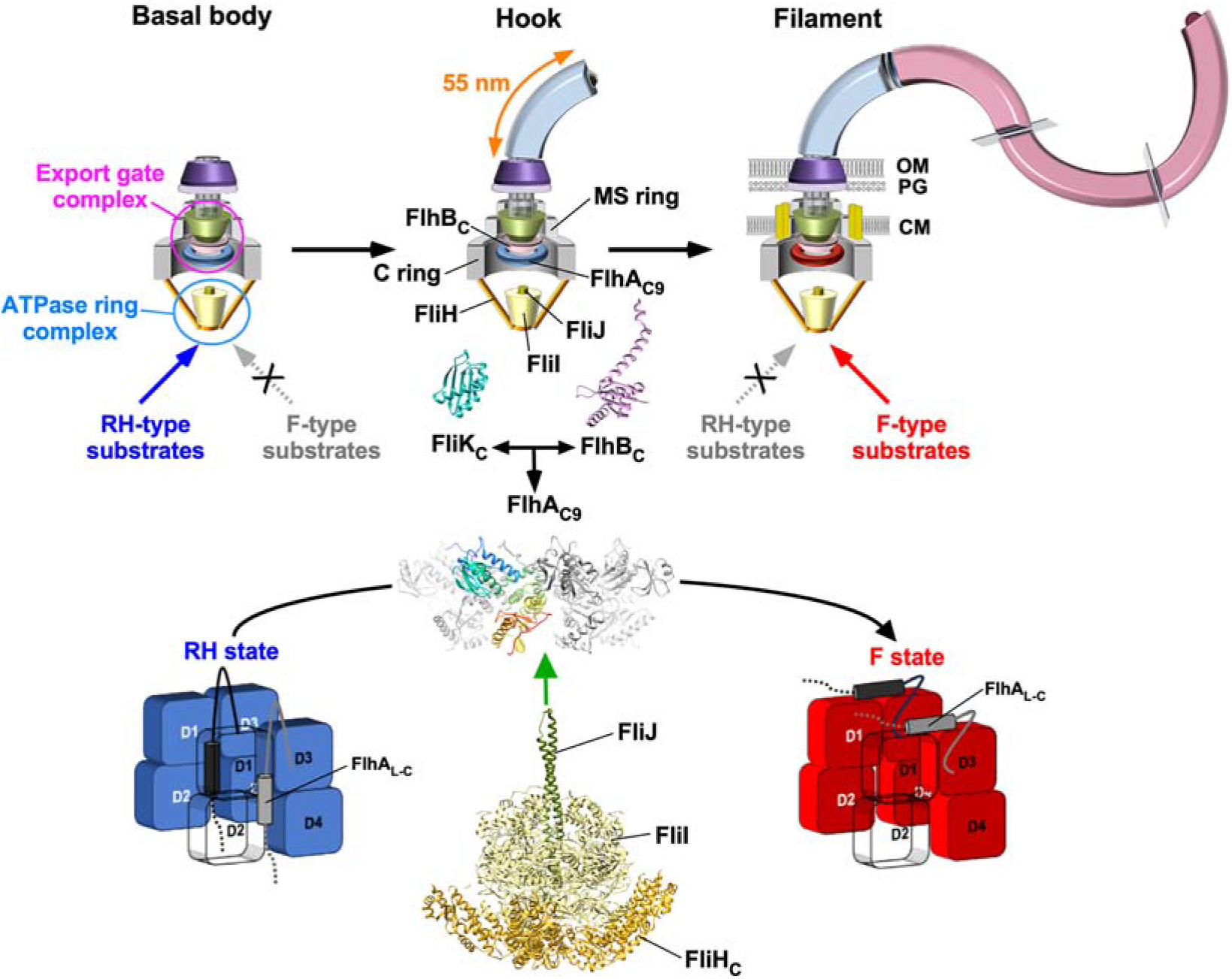
Model for substrate specificity switching of the flagellar type III secretion system. Flagellar assembly begins with the basal body, followed by the hook and finally the filament. The flagellar type III secretion system (fT3SS) transports 14 different proteins from the cytoplasm during flagellar assembly. These 14 proteins are classified into two export classes: One is the RH-type substrates needed for the structure and assembly of the rod and hook, and the other is the F-type substrates responsible for filament formation. The fT3SS consists of a transmembrane export gate complex and a cytoplasmic ATPase ring complex consisting of FliH, FliI, and FliJ. The export gate complex is located within the MS ring. The C-terminal cytoplasmic domains of FlhA (FlhA_C_) and FlhB (FlhB_C_) project into the central cavity of the C ring and are directly involved in substrate specificity switching of the fT3SS from the RH-type to the F-type upon hook completion. The ATPase ring complex associates with the C ring. During hook-basal body assembly, the fT3SS transports the RH-type substrates but not the F-type substrates. When the hook length reaches about 55 nm, the C-terminal domain of FliK (FliK_C_) binds to FlhB_C_, followed by a structural transition of the FlhA_C_ ring (FlhA_C9_) from the RH state to the F state, allowing the fT3SS to terminate RH-type protein export and initiate F-type protein export. The present study establishes that the cytoplasmic ATPase complex is required for the FlhA_C_ ring to efficiently switch its conformation from the RH state to the F state. Atomic models of FliK_C_ (PDB ID: 2RRL), FlhB_C_ (PDB ID: 3B0Z), FlhA_C_ (PDB ID: 3A5I), the FliH_C2_-FliI complex (PDB ID: 5B0O), and FliJ (PDB ID: 3AJW) are shown in Cα ribbon representation.

The *Salmonella* fT3SS transports 14 different proteins from the cytoplasm during flagellar formation. These 14 proteins are classified into two distinct types, rod-hook-type (hereafter referred to as RH-type) and filament-type (hereafter referred to as F-type) classes^14^. During hook-basal body (HBB) assembly, the fT3SS acts specially on the RH-type substrates needed for assembly of the rod and hook. Four F-type substrates, FlgK, FlgL, FlgM, and FliD, are expressed during HBB assembly, but are not transported by the fT3SS^15^. Once the HBB is complete, the fT3SS switches substrate specificity from the RH-type to the F-type and hence acts specially on the F-type substrates to build the filament at the hook tip (Fig. 1). To coordinate flagellar protein export with assembly, the fT3SS uses a secreted molecular ruler named FliK not only to measure the length of the hook but also to catalyze the substrate specificity switching of the fT3SS when the hook length reaches about 55 nm^16–20^. The export switch of the fT3SS consists of the C-terminal cytoplasmic domains of FlhA (FlhA_C_) and FlhB (FlhB_C_)^21,22^. Genetic analyses combined with photo-crosslinking experiments have suggested that a direct interaction between FliK and FlhB_C_ causes a conformational change in FlhB_C_, followed by conformational rearrangements of FlhA_C_ that terminates hook assembly and initiates filament assembly^23–25^. Therefore, in *Salmonella* strains with loss-of-function of FliK or specific amino acid substitutions in FlhA_C_ or FlhB_C_, the substrate recognition mode of the fT3SS remains in the RH state, resulting in unusually elongated hooks named polyhooks.

FlhA_C_ consists of four domains, D1, D2, D3, and D4, and a flexible linker (FlhA_L_) connecting FlhA_C_ with the N-terminal transmembrane domain of FlhA (Supplementary Fig. 1a)^26^. FlhA_C_ forms a nonameric ring through intermolecular interactions between domains D1 and D1 and between domains D1 and D3 (Fig. 1)^27^. The highly conserved Asp-456 and Thr-490 residues are very close to each other and are located within a hydrophobic dimple formed by the relatively well-conserved Leu-438, Ile-440, Pro-442, Phe-459, Leu-461, Val-482, and Val-487 residues (Supplementary Fig. 1a). This conserved dimple, located at the interface between domains D1 and D2, is directly involved in substrate recognition^28–30^. During hook assembly, the C-terminal region of FlhA_L_ (FlhA_L-C_) including the Glu-351, Trp-354, and Asp-356 residues binds to the conserved hydrophobic dimple, not only facilitating the export of the hook protein (FlgE) but also suppressing the interaction of FlhA_C_ with flagellar export chaperones (FlgN, FliS, FliT) in complex with their cognate F-type substrates^31^. Upon completion of HBB assembly, FlhA_L-C_ is detached from the conserved dimple and binds to the D1 and D3 domains of its neighboring FlhA_C_ subunit in the ring^22,32^. As a result, the fT3SS switches the substrate recognition mode from the RH state to the F state (Fig. 1). High-speed atomic force microscopy (HS-AFM) has shown that the W354A, E351A/D356A, and E351/W354A/D356A mutations in FlhA_L-C_ inhibits FlhA_C_ ring formation. Furthermore, these mutations reduce the binding affinity of FlhA_C_ for flagellar export chaperones in complex with their cognate F-type substrates, thereby preventing F-type protein export^22^. These suggest that FlhA_L-C_ acts as a structural switch that induces the structural transition of the FlhA_C_ ring from the RH state to the F state. However, it remains unknown how it occurs upon hook completion.

Gly-368 of *Salmonella* FlhA is located within the highly conserved GYXLI motif in domain D1 of FlhA_C_ and is important for dynamic domain motions required for efficient and rapid protein export by the fT3SS (Supplementary Fig. 1a)^33,34^. The temperature-sensitive *flhA(G368C)* mutant cultured at 42°C produces no flagella but at 30°C produces almost the same number of flagella as wild-type cells^35^, and its average hook length is comparable to that of the wild-type^33^. In contrast, the Δ*fliH-fliI flhB(P28T)* strain (hereafter referred to as ΔHI-B*), which produces a few flagella even in the absence of FliH and FliI^5^, produces no filaments even at 30°C when the *flhA(G368C)* mutation is added^33^. Furthermore, the average hook length of the ΔHI-B* *flhA(G368C)* strain is longer than that of the ΔHI-B*strain despite the presence of FliK^33^, leading to a plausible hypothesis that FliH and FliI may also be required for a conformational change of the conserved GYXLI motif to induce the transition of the FlhA_C_ ring structure from the RH state to the F state.

Here, to clarify this hypothesis, we focused on the conserved GYXLI motif of FlhA_C_. We performed genetic analyses of the Y369A/R370A/L371A/I372A (hereafter referred to as AAAA) and Y369G/R370G/L371G/I372G (hereafter referred to as GGGG) mutants and provide evidence suggesting that FliH and FliI are not only required for FliK-dependent remodeling of the FlhA_C_ ring structure through conformational changes in the conserved GYXLI motif, but also for FlhA_C_ to exert its proofreading function to correct substrate recognition errors during flagellar formation.

## Results

### Effect of the AAAA and GGGG mutations on flagellar protein export

It has been reported that the AAAA and GGGG mutations inhibit flagella-driven motility in soft agar (Fig.2a, left panel)^34^. To clarify how these two mutations affect the flagellar assembly process, we analyzed the secretion levels of RH-type proteins, such as FlgD, FlgE, and FliK, and F-type proteins, such as FlgM, FlgK, FlgL, and FliC, by immunoblotting (Supplementary Fig. 1b). The levels of FlgD (1st row), FlgE (2nd row), and FliK (3rd row) secreted by the AAAA and GGGG mutants were higher than the wild-type levels. In contrast to the RH-type substrates, the AAAA mutation reduced the secretion levels of FlgM (4th row), FlgK (5th row), FlgL (6th row), and FliC (7th row) whereas the GGGG mutation inhibited the secretion of these four F-type proteins. Consistently, the AAAA mutant produced short flagellar filaments whereas the GGGG mutant produced no filaments (Supplementary Fig. 2). These results strongly suggest that these two mutations affect substrate specificity switching of the fT3SS from the RH-type to the F-type.

To examine whether the AAAA and GGGG mutants produce polyhooks, HBBs were purified from these two mutants, and their hook length was measured (Supplementary Fig. 1c). The hook lengths of the AAAA and GGGG mutants were 225.7 ± 176.8 (mean ± SD) nm (N = 300) and 471.1 ± 285.1 nm (N = 300), respectively, indicating that the fT3SS with the AAAA or GGGG mutation cannot terminate hook assembly at an appropriate timing of hook assembly or at all even when both FliK and FlhB are intact. Therefore, we conclude that the substrate specificity of fT3SS is determined by the conformational state of the FlhA_C_ ring structure and that these two mutations severely impair or inhibit the conformational transition of the FlhA_C_ ring from the RH state to the F state. Because conformational changes in the GYXLI motif of FlhA_C_ cause a reorganization of the hydrophobic side-chain interaction network throughout the entire FlhA_C_ structure, allowing FlhA_C_ to take different conformations ^34^, a specific conformational change of the GYXLI motif would be necessary for the efficient and robust structural transition of the FlhA_C_ ring from the RH state to the F state upon hook completion.

### Isolation of up-motile mutants from the AAAA and GGGG mutants

To clarify how the AAAA and GGGG mutations inhibit substrate specificity switching of the fT3SS, we isolated seven and three up-motile mutants from the AAAA and GGGG mutants, respectively. Improvements in motility of the up-motile mutants are shown in the left panel of Fig. 2a and Fig. 3a. DNA sequencing of the seven up-motile mutants isolated from the AAAA mutant identified two missense mutations, A372V (isolated three times) and A372T in FlhA_C_ (Fig. 2b), and two missense mutations, Q338R and A405V (isolated twice) in FliK (Fig. 3b). The three up-motile mutants isolated from the GGGG mutant revealed that all suppressor mutations were G372V missense mutations in FlhA_C_ (Fig. 2b). All the intragenic suppressor mutations share a common feature that the change of residue occurred at position 4 of the four mutated residues in the parent *flhA* mutants, such as from AAAA to AAAV or AAAT and from GGGG to GGGV.

**Fig. 2.**
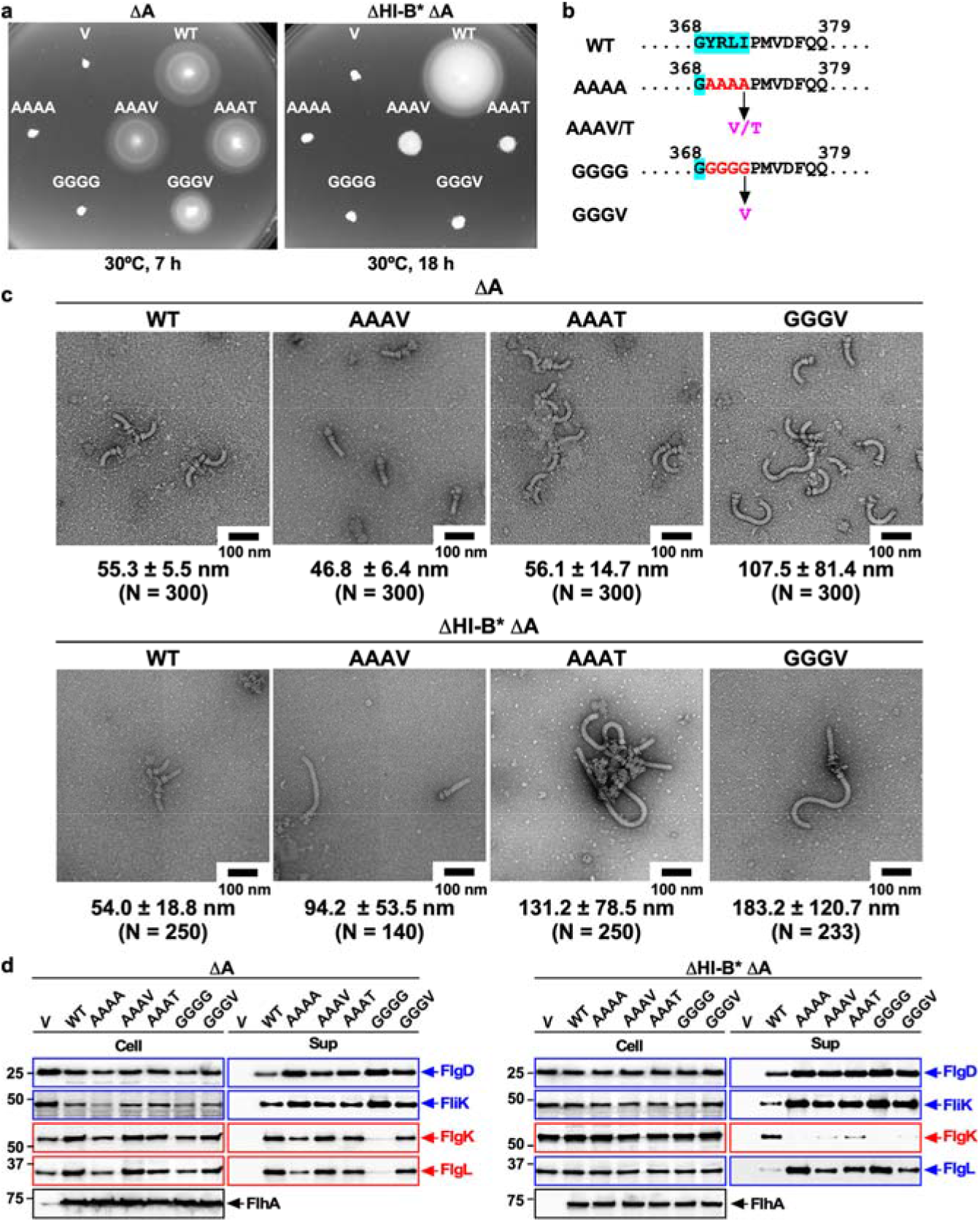
Effect of mutations in the conserved GYXLI motif of FlhA on flagellar protein export and assembly in the presence and absence of FliH and FliI. **(a)** Motility of the *Salmonella* NH001 (Δ*flhA,* indicated as ΔA) or NH003 [Δ*fliH-fliI flhB(P28T)* Δ*flhA*, indicated as ΔHI-B* ΔA] strain transformed with pTrc99AFF4 (indicated as V), pMM130 (indicated as WT), pMKM130-A4 (indicated as AAAA), pMKM130-A3V (indicated as AAAV), pMKM130-A3T (indicated as AAAT), pMKM130-G4 (indicated as GGGG), or pMKM130-G3V (indicated as GGGV) in the presence (left panel) and absence (right panel) of FliH and FliI. Soft tryptone agar plates were incubated at 30°C for 7 hours (left panel) or 18 hours (right panel). **(b)** Location of intragenic suppressor mutations isolated from the AAAA and GGGG mutants. The conserved GYXLI motif of FlhA is highlighted in cyan. The intragenic A372V or A372T suppressor mutation isolated from the AAAA mutant is the change of alanine at position 4 in the AAAA sequence to valine or threonine, respectively, and the intragenic G372V suppressor mutation is the change of glycine at position 4 in the GGGG sequence to valine. **(c)** Electron micrographs of hook-basal bodies isolated from the above transformants. The average hook length and standard deviations are shown. N indices the number of hook-basal bodies and polyhook-basal bodies that were measured. **(d)** Secretion analysis of FlgD, FliK, FlgK, and FlgL by immunoblotting. Whole cell proteins (Cell) and culture supernatants (Sup) were prepared from the above transformants. A 3 μl solution of each sample normalized to an optical density of OD_600_ was subjected to SDS-PAGE and analyzed by immunoblotting using polyclonal anti-FlgD (1st row), anti-FliK (2nd row), anti-FlgK (3rd row), anti-FlgL (4th row), or anti-FlhA_C_ (5th row) antibody. RH-type and F-type substrates are highlighted in blue and red, respectively. Molecular mass markers (kDa) are shown on the left.

**Fig. 3.**
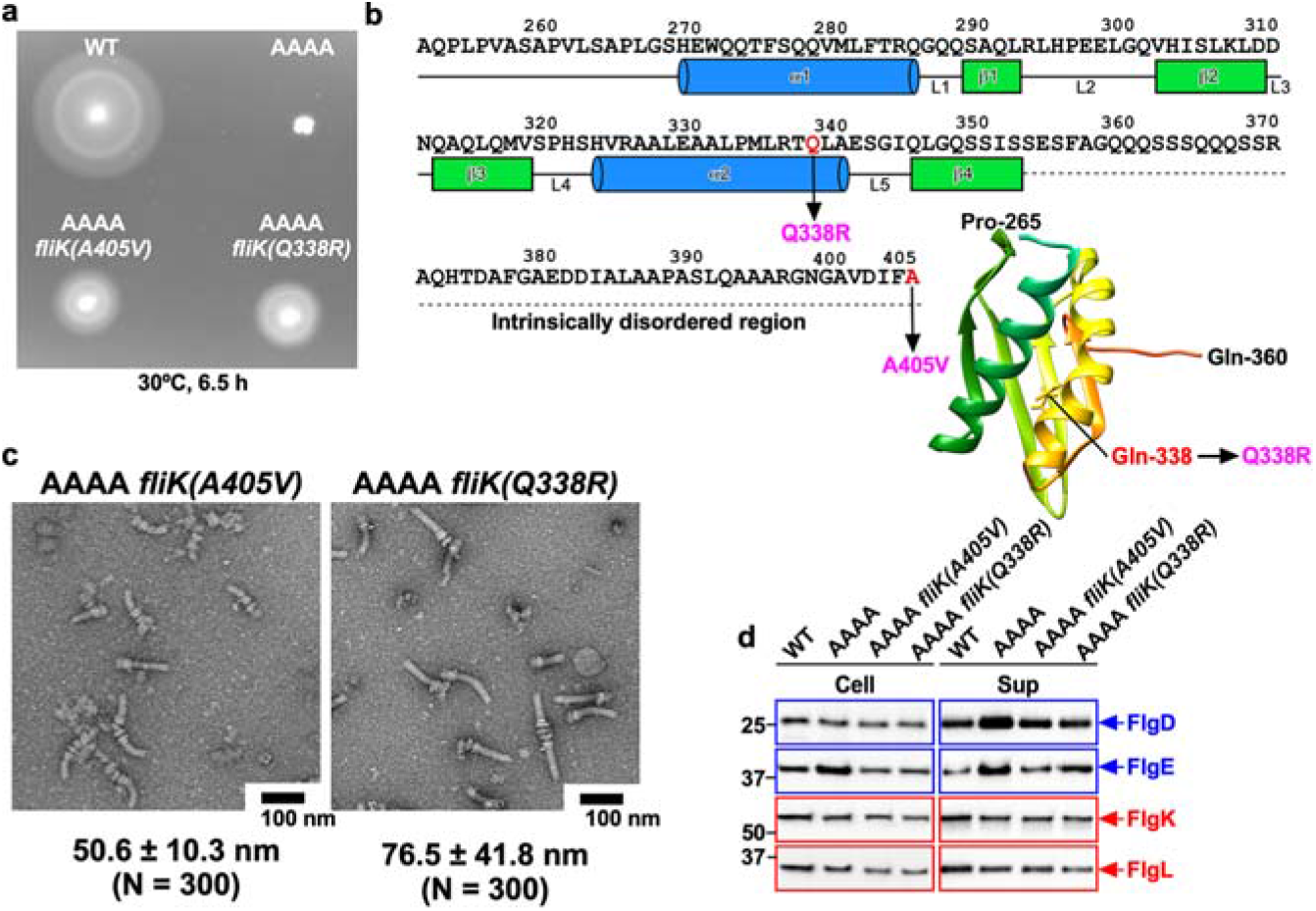
Isolation of extragenic suppressor mutants from the AAAA mutant. **(a)** Motility of NH001 carrying pMM130 (WT), MMA130A4 (AAAA), MMA130A4-3 [AAAA *fliK(A405V)*], and MMA130A4-10 [AAAA *fliK(Q338R)*] in soft agar. Plates were incubated at 30°C for 6.5 hours. **(b)** Location of extragenic suppressor mutations in the C-terminal domain of FliK (FliK_C_). The compactly folded core domain of FliK_C_ (PDB ID: 2RRL) consists of residues 268–352 and is directly involved in substrate specificity switching of the fT3SS from the RH-type to the F-type. The secondary structures are shown below the amino acid sequence of FliK. Residues of 353–405 are intrinsically disordered in solution. The Cα backbone is color-coded from green to orange, going through the rainbow colors from the N- to C-terminus. Extragenic suppressor mutations are highlighted in magenta. **(c)** Electron micrographs of hook-basal bodies isolated from the above strains. The average hook length and standard deviations are shown. N indices the number of hook-basal bodies that were measured. **(d)** Secretion analysis of FlgD, FlgE, FlgK, and FlgL by immunoblotting. Whole cell proteins (Cell) and culture supernatants (Sup) were prepared from the above strains. A 3 μl solution of each sample normalized to an optical density of OD_600_was subjected to SDS-PAGE and analyzed by immunoblotting using polyclonal anti-FlgD (1st row), anti-FlgE (2nd row), anti-FlgK (3rd row), or anti-FlgL (4th row) antibody. RH-type and F-type substrates are highlighted in blue and red, respectively. Molecular mass markers (kDa) are shown on the left.

### Characterization of intragenic AAAV, AAAT, and GGGV suppressor mutants

To examine whether intragenic suppressor mutations shortened the length of polyhooks produced by the AAAA and GGGG mutants, HBBs were purified from the AAAV, AAAT, and GGGV mutants, and their hook length was measured (Fig 2c, upper panel). The hook lengths of the AAAV, AAAT, and GGGV mutants were 46.8 ± 6.4 nm (N = 300), 56.1 ± 14.7 nm (N = 300), and 107.5 ± 81.4 nm (N = 300), respectively, which are much shorter than 225.7 ± 176.8 nm (N = 300) of the AAAA mutant cells and 471.1 ± 285.1 nm (N = 300) of the GGGG mutant cells and are much closer to 55.3 ± 5.5 nm (N =300) of wild-type cells. This indicates that these suppressor mutations considerably shortened the polyhook length but did not recover the precise control of the hook length of the wild type. Consistently, these suppressor mutations reduced the secretion levels of FlgD, FlgE, and FliK compared to the AAAA and GGGG mutants (Fig. 2d, left panel, 1st and 2nd rows and Supplementary Fig. 3, left panel, 1st row). A loss-of-function of FliK caused complete inhibition in the motility of these three suppressor mutants in soft agar (Supplementary Fig. 4a) by producing polyhooks without filament attached (Supplementary Fig. 4b). These results suggest that the fT3SS with the AAAV, AAAT, or GGGV mutation can receive the hook length signal from the FliK ruler protein and terminate the export of RH-type substrates at a more appropriate timing of hook assembly compared to their parental mutant strains.

To investigate whether these intragenic suppressor mutations restore the export of F-type substrates to the wild-type levels, we analyzed the secretion levels of FlgK, FlgL, FliC, and FliD by immunoblotting (Fig. 2d, left panel, 3rd and 4th rows and Supplementary Fig. 3, left panel, 2nd and 3rd rows). The amounts of FlgK, FlgL, FliC, and FliD secreted by the AAAV and AAAT suppressor mutants were greater than those seen in the AAAA mutant and similar to the wild-type levels. Consistently, they produced longer flagellar filaments than the AAAA mutant (Supplementary Fig. 2). The secretion levels of F-type substrates by the GGGV suppressor mutant were also much higher than those by the GGGG mutant, but lower than the wild-type levels (Fig. 2d, left panel, 3rd and 4th rows and Supplementary Fig. 3, left panel, 2nd and 3rd rows). Consistent with this, the GGGV suppressor mutant produced shorter flagellar filaments than wild-type cells (Supplementary Fig. 2). Because these F-type substrates require their cognate export chaperones for efficient docking to the FlhA_C_ ring for export^28,29^, these results suggest that, compared to FlhA_C_ with the AAAA or GGGG mutation, FlhA_C_ with the AAAV, AAAT, or GGGV mutation can make a more appropriate chaperone binding site in the well-conserved hydrophobic dimple of FlhA_C_once HBB assembly is complete. Therefore, we conclude that the conformational change of the GYXLI motif of FlhA_C_ is necessary not only for efficient transition of the FlhA_C_ ring from the RH state to the F state but also for the formation of appropriate chaperone binding sites in the ring.

### Characterization of extragenic suppressor mutants isolated from the AAAA mutant

The extragenic suppressor mutations isolated from the AAAA mutant, *fliK(A405V)* and *fliK(Q338R)*, are located within FliK_C_, which directly binds to FlhB_C_ and catalyzes substrate specificity switching of the fT3SS from the RH-type to the F-type (Fig. 3b). To investigate how these two extragenic suppressor mutations improve flagella-driven motility of the AAAA mutant in soft agar (Fig. 3a), HBBs were purified, and their hook length was measured. The hook lengths of the AAAA *fliK(A405V)* and AAAA *fliK(Q338R)* mutants were 50.6 ± 10.3 nm (N = 300) and 76.5 ± 41.8 nm (N = 300), respectively (Fig. 3c), compared to 225.7 ± 176.8 nm (N = 300) of the AAAA mutant. This indicates that the fT3SS with the AAAA mutation can receive the hook length signal more efficiently from FliK with the A405V or Q338R mutation than that from wild-type FliK and thus can stop the export of RH-type proteins at a more appropriate timing of hook assembly compared to the AAAA mutant. In agreement with this, these two *fliK* mutations reduced the secretion levels of FlgD and FlgE (Fig. 3d, 1st and 2nd rows). To investigate the export switching efficiency of these extragenic suppressor mutants, we analyzed the secretion levels of FlgK and FlgL by immunoblotting (Fig. 3d, 3rd and 4th rows). The levels of FlgK and FlgL secreted by the AAAA *fliK(A405V)* and AAAA *fliK(Q338R)* mutants were essentially the same as those seen in the AAAA mutant. Consistently, they produced short flagellar filaments in a way like the AAAA mutant (Supplementary Fig. 2). These observations suggest that the termination of RH-type protein export and activation of F-type protein export are independent processes that are not tightly coupled with each other and can be interpreted as that FlhA_C_ with the AAAA mutation in the ring may efficiently shift its conformation from the RH state to the F state to terminate hook assembly through the action of FliK with the extragenic suppressor mutations but cannot make an appropriate chaperone binding site.

To test whether these second site *fliK* mutations by themselves affect the export switching function of the fT3SS, the *fliK* mutations were transferred by P22 phage-mediated transduction into TH8426 (Δ*fliK*) to construct strains containing only either *fliK(A405V)* or *fliK(Q338R)* mutation. Motility of the *fliK(A405V)* and *fliK(Q338R)* mutants were almost the same as that of wild-type cells (Supplementary Fig. 5), indicating that these *fliK* mutations display no significant motility phenotype. Therefore, we propose that these *fliK* mutations may induce a conformational change in FliK_C_ to allow its stronger action on FlhB_C_, whereby inducing a conformational change of the FlhA_C_ ring with the AAAA mutation for more efficient termination of RH-type protein export.

### Effect of mutations in the GYXLI motif on FlhA_C_ ring formation

A combination of HS-AFM and structure-based mutational analysis has shown that the interactions of FlhA_L-C_ with the D1 and D3 domains of its closest FlhA_C_ subunit are important not only for FlhA_C_ ring formation on mica surfaces but also for substrate specificity switching of the fT3SS from the RH-type to the F-type, suggesting that the FlhA_C_ ring observed by HS-AFM reflects an F-type ring structure^22^. To obtain direct evidence that the GYXLI motif of FlhA is involved in the structural transition of FlhA_C_ from the RH state to the F state, we purified the N-terminally His-tagged FlhA_C-AAAA_ and FlhA_C-GGGG_ monomers by size exclusion chromatography with a Superdex 75 column HR 10/30 column (Supplementary Fig. 6a) and analyzed their ring forming ability by HS-AFM (Fig. 4). FlhA_C-AAAA_ formed the ring structure like wild-type FlhA_C_ whereas FlhA_C-GGGG_ did not. This indicates that the GGGG mutation inhibits the interaction of FlhA_L-C_ with its closest FlhA_C_ subunit while the AAAA mutation does not. Because far-UV CD measurements revealed that the AAAA and GGGG mutation did not affect the secondary structures of FlhA_C_ (Supplementary Fig. 6b), we conclude that a conformational change of the GYXLI motif induces a structural transition of the FlhA_C_ ring from a relatively unstable RH-state ring to a more stable F-state ring to terminate hook assembly.

**Fig. 4.**
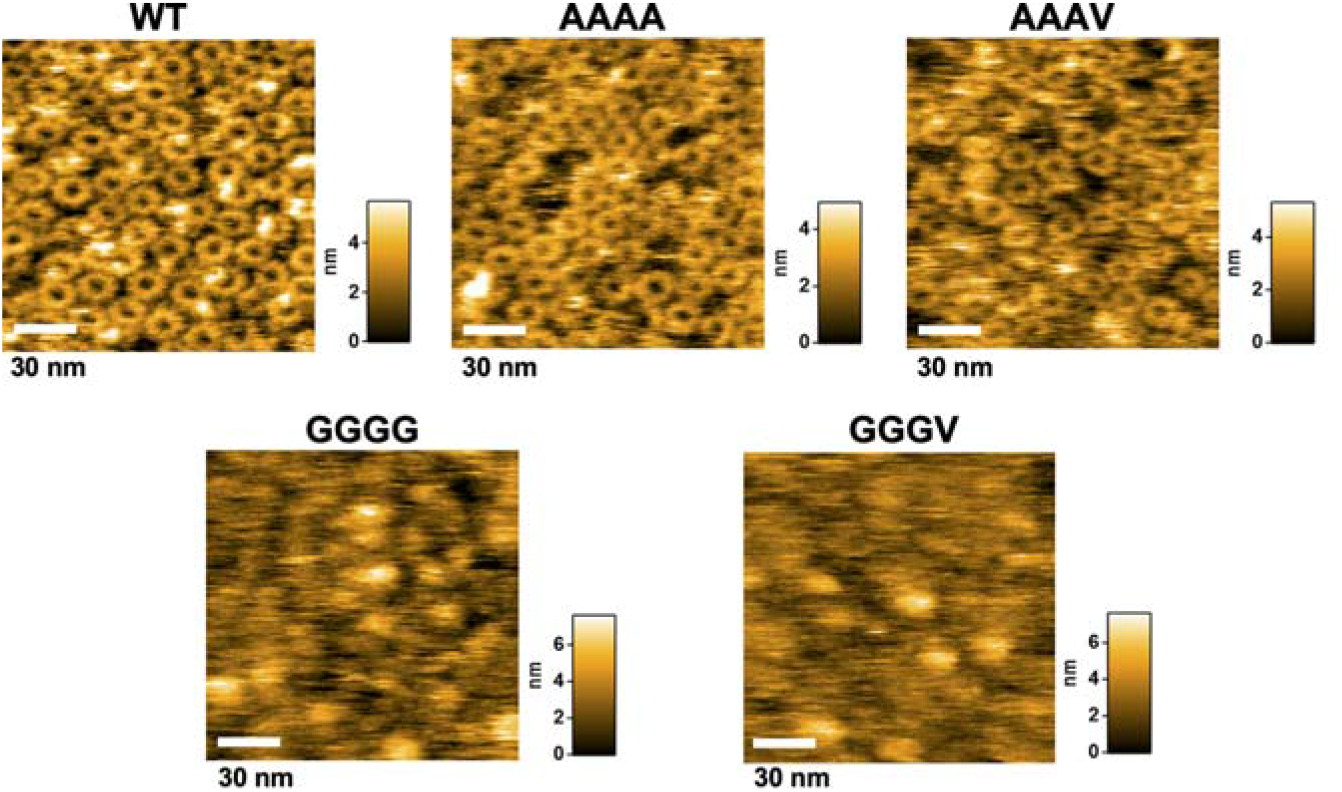
Effect of mutations in the GYXLI motif on FlhA_C_ ring formation. Typical HS-AFM images of His-FlhA_C_ (WT) and its mutant variants with either AAAA, AAAV, GGGG, or GGGV mutation placed on mica surface in a buffer at a protein concentration of 2 μM. All images were recorded at 200 ms/frame in a scanning area of 100 × 100 nm^2^ with 150 ×150 pixels. Color bar on the right of each image indicates a range of particle heights (nm).

We next investigated whether the intragenic suppressor mutations affect FlhA_C_ ring formation. FlhA_C-AAAV_ formed the ring structure as FlhA_C-AAAA_ but FlhA_C-GGGV_ failed to form the ring structure as FlhA_C-GGGG_. This was rather surprising because the fT3SS with FlhA_C-GGGV_ can switch its substrate specificity from the RH-type to the F-type to a significant degree (Fig. 2). So, we hypothesized that the GGGG and GGGV mutations both lock FlhA_C_ in the RH state but FlhA_C_ with the GGGV mutation can make a structural transition to the F state may be with a support of other proteins when it receives the hook length signal from FliK.

### Effect of mutations in the GYXLI motif on the interaction of FlhA_C_ with the FlgN-FlgK chaperone-substrate complex

The FlgN-FlgK/L, FliS-FliC, and FliT-FliD chaperone-substrate complexes directly bind to a well-conserved hydrophobic dimple located at an interface between domains D1 and D2 of FlhA_C_, allowing the fT3SS to efficiently transport F-type proteins into the central channel of the growing flagellar structure and to its distal end for assembly^28–30^. Because the AAAA mutation reduced the secretion levels of F-type protein export by the fT3SS whereas its intragenic AAAV suppressor mutation restored the secretion levels to the wild-type level (Fig. 2c, left panel), we investigated whether these mutations affect the interaction of FlhA_C_ with the FlgN-FlgK chaperone-substrate complex by GST affinity chromatography (Fig. 5a). Wild-type FlhA_C_ co-purified with GST-FlgN in complex with FlgK (1st row), in agreement with previous reports^28,29^. In contrast, only a very small amount of FlhA_C-AAAA_ co-purified with the GST-FlgN/FlgK complex (2nd row), and its intragenic suppressor mutation increased the binding affinity for the FlgN/FlgK complex although not to the wild-type level (3rd row). Because neither the AAAA nor AAAV mutation inhibited FlhA_C_ ring formation (Fig. 4), we conclude that a proper conformational change of the GYXLI motif of FlhA_C_ is also required for the formation of an appropriate chaperone binding site in the hydrophobic dimple of FlhA_C_ after FlhA_L-C_ dissociates from the hydrophobic dimple and binds to the D1 and D2 domains of the closest FlhA_C_subunit in the FlhA_C_ ring.

**Fig. 5.**
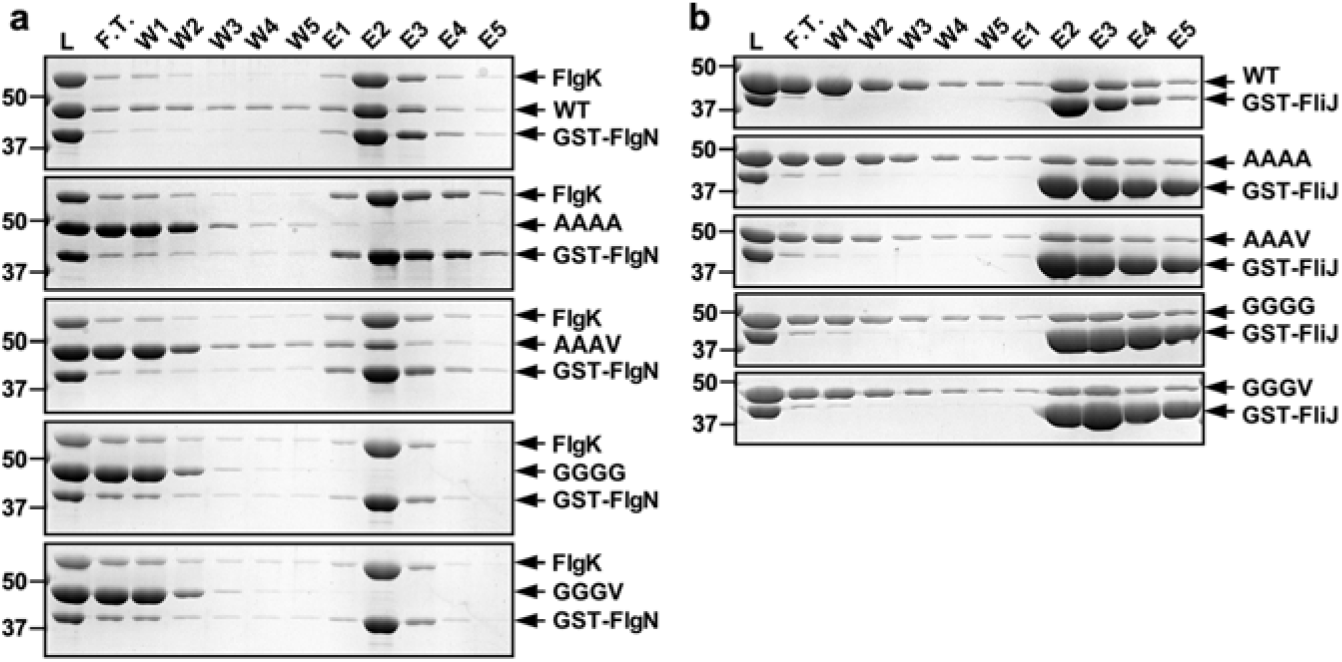
Effect of mutations in the GYXLI motif on the interaction of FlhA_C_ with the FlgN-FlgK chaperone-substrate complex and FliJ. Mixtures (L) of purified His-FlhA_C_ (WT, 1st row), His-FlhA_C-AAAA_ (AAAA, 2nd row), His-FlhA_C-AAAV_ (AAAV, 3rd row), His-FlhA_C-GGGG_ (GGGG, 4th row), or His-FlhA_C-GGGV_ (GGGV, 5th row) with GST-FlgN in complex with FlgK **(a)** or GST-FliJ **(b)** were dialyzed overnight against PBS, followed by GST affinity chromatography. Flow through fraction (F.T.), wash fractions (W) and elution fractions (E) were analyzed by CBB staining. Molecular mass markers (kDa) are shown on the left.

Because the GGGG and GGGV mutations both inhibited FlhA_C_ ring formation (Fig. 4), we investigated whether they also prevent the FlgN-FlgK complex from binding to FlhA_C_. FlhA_C_ with the GGGG or GGGV mutation did not co-purified with the GST-FlgN/FlgK complex at all (Fig. 5a, 4th and 5th rows). Therefore, we conclude that the GGGG and GGGV mutations stabilize the FlhA_C_ conformation in the RH state, thereby inhibiting the interaction of FlhA_C_ with the FlgN-FlgK complex *in vitro*.

### Effect of mutations in the GYXLI motif on the interaction of FlhA_C_ with FliJ

The W354A and E351A/D356A mutations in FlhA_L-C_ not only inhibit FlhA_C_ ring formation but also reduce the binding affinity of FlhA_C_ for flagellar export chaperones in complex with their cognate F-type substrates, and these two mutations also reduce the binding affinity of FlhA_C_ for FliJ^22,31^, leading a plausible hypothesis that the FlhA_C_-FliJ interaction may be required for efficient transition of the FlhA_C_ ring structure from the RH state to the F state. Therefore, we analyzed the FlhA_C_-FliJ interaction by GST affinity chromatography (Fig. 5b). FlhA_C_ with the AAAA, AAAV, GGGG, or GGGV mutation co-purified with GST-FliJ as wild-type FlhA_C_, indicating that these mutations do not reduce the binding affinity of FlhA_C_ for FliJ. Therefore, we conclude that the GYXLI motif of FlhA_C_ is not involved in the interaction with FliJ.

### Effect of FliH and FliI deletion on the export switching function of FlhA with the AAAV, AAAT or GGGV mutation

FliH and FliI, components of the fT3SS ATPase complex, are thought not only to ensure efficient and robust energy coupling of flagellar protein export but also to contribute to hierarchical protein targeting of export substrates and chaperone-substrate complexes to the FlhA_C_ ring^36–38^. Because the AAAV and GGGV mutations both reduced the binding affinity of FlhA_C_ for the FlgN-FlgK chaperone-substrate complex (Fig. 5a), we investigated if FlhA with either AAAV, AAAT, or GGGV mutation requires the support of FliH and FliI to fully exert its export function. Because neither flagella-driven motility in soft agar nor flagellar protein export by the fT3SS was affected by the *flhB(P28T)* (hereafter referred to as B*) mutation alone (Supplementary Fig. 7), we analyzed the effect of the AAAV, AAAT, or GGGV mutation on the export switching function of FlhA in the ΔHI-B* mutant background. Unlike in the presence of FliH and FliI with or without the B* mutation, the motility of the ΔHI-B* AAAV, ΔHI-B* AAAT, and ΔHI-B* GGGV mutants was much worse than that of the ΔHI-B* mutant (Fig. 2a, right panel). Consistently, the secretion levels of F-type substrates such as FlgK, FliC, and FliD were lower in these three mutants than in the ΔHI-B* strain (Fig. 2d, right panel, 3rd row and Supplementary Fig. 3, right panel, 2nd and 3rd rows). Interestingly, much longer polyhooks were frequently observed in these three mutants compared to the AAAV, AAAT, and GGGV mutants, respectively (Fig. 2c, lower panel). Therefore, we quantitatively measured the polyhook length of the three mutants. The hook length of the ΔHI-B* strain was 54.0 ± 18.8 nm (N = 250), showing a much broader length distribution compared to the wild-type, in agreement with a previous report^38^. The polyhook lengths of the ΔHI-B* AAAV, ΔHI-B* AAAT, and ΔHI-B* GGGV mutants were 94.2 ± 53.5 nm (N = 140), 131.2 ± 78.5 nm (N = 250), and 181.2 ± 120.7 nm (N = 233), respectively, compared to 46.8 ± 6.4 nm (N = 300) for the AAAV mutant, 56.1 ± 14.7 nm (N = 300) for the AAAT mutant, and 107.5 ± 81.4 nm (N = 300) for the GGGV mutant cells. Consistently, the amounts of RH-type substrates such as FlgD, FlgE, and FliK secreted from the ΔHI-B* AAAV, ΔHI-B* AAAT, and ΔHI-B* GGGV mutants were higher than those of the ΔHI-B* mutant (Fig. 2d, right panel, 1st and 2nd rows and Supplementary Fig. 3, right panel, 1st row). These results suggest that the FlhA_C_ ring with the AAAV, AAAT or GGGV mutation requires the support of FliH and FliI to efficiently dissociate FlhA_L-C_ from the hydrophobic dimple of FlhA_C_ when it receives the hook length signal from FliK.

### Effect of the AAAA and GGGG mutations on FlgL secretion in the presence and absence of FliH and FliI

We found a peculiar change in the secretion level of FlgL, one of the F-type substrates, by the presence and absence of FliH and FliI (Fig. 2d, 4th rows and Supplementary Fig 7b, 5th row). In the presence of FliH and FliI, the amount of FlgL secreted from the AAAA and GGGG mutants, with or without the B* mutation, was less than those of the wild-type strain and their intragenic suppressor mutants, as was the case for other F-type proteins. However, in the absence of FliH and FliI, the amount of FlgL secreted from the AAAA and GGGG mutants was higher than those secreted from the wild-type and their suppressor mutants, as was the case for the RH-type proteins. Because the AAAA and GGGG mutations impair substrate specificity switching of the fT3SS from the RH-type to the F-type at an appropriate timing of hook assembly even in the presence of FliH and FliI (Supplementary Fig. 1), we suggest that the fT3SS with the AAAA or GGGG mutation recognizes FlgL as an RH-type substrate rather than an F-type substrate in the absence of FliH and FliI whereas it recognizes FlgL properly as an F-type substrate in the presence of FliH and FliI.

Both FlgK and FlgL require the FlgN chaperone for efficient binding to FlhA_C_, allowing these two proteins to be efficiently unfolded and transported by the fT3SS. So, when the interaction between FlgN and FlhA is impaired, the secretion levels of FlgK and FlgL are significantly reduced^28,29^. To investigate whether FlgL secretion by the ΔHI-B* AAAA and ΔHI-B* GGGG mutants is dependent on FlgN, we introduced a Δ*flgN*::*tetRA* allele into the ΔHI-B* ΔA, ΔHI-B*, ΔHI-B* AAAA and ΔHI-B* GGGG mutants by P22-mediated transduction and analyzed the secretion level of FlgL by immunoblotting. FlgN has been reported to be essential for the export of both RH-type and F-type substrates because it also acts as an activator of the transmembrane export gate complex of the fT3SS when the cytoplasmic ATPase complex is dysfunctional^10^. Therefore, we also measured the secretion level of FlgD (Supplementary Fig. 8). As expected, neither FlgD nor FlgL was secreted from the ΔHI-B* cells containing the Δ*flgN*::*tetRA* allele. The AAAA and GGGG mutations overcame the effect of FlgN deletion on flagellar protein export, thereby allowing both FlgD and FlgL to be secreted extracellularly even in the absence of FliH and FliI. This indicates that the fT3SS with the AAAA or GGGG mutation does not require FlgN for FlgL secretion in the absence of FliH and FliI. Since the *flhA(D456V)* and *flhA(T490M)* mutations in the conserved hydrophobic dimple of FlhA_C_ have been shown to be able to bypass the FlgN defect to a significant degree^10,28^, the AAAA and GGGG mutations seem to induce a required conformational change in the conserved dimple of FlhA_C_, allowing the transmembrane export gate complex to become an active protein transporter.

## Discussion

When the hook length reaches about 55 nm in *Salmonella*, FliK catalyzes substrate specificity switching of the fT3SS from the RH-type to the F-type through a direct interaction between FliK_C_ and FlhB_C_, thereby terminating hook assembly and initiating filament formation. The FlhA_C_ ring takes at least two distinct conformational states, RH and F. However, it remained unclear how the FlhA_C_ ring switches its conformation from the RH state to the F state upon hook completion. To answer this question, we focused on the highly conserved GYXLI motif of FlhA_C_, which acts as a structural switch to facilitate cyclic open-close domain motions of FlhA_C_ through periodically remodeling its hydrophobic side-chain interaction networks^34^ and discovered that the GYXLI motif, together with the fT3SS ATPase complex, is directly involved not only in the substrate specificity switching but also in the proofreading function of the fT3SS, thereby bringing strict order to flagellar protein export for efficient assembly of the flagellum.

During HBB assembly, FlhA_L-C_ binds to a well-conserved hydrophobic dimple of FlhA_C_, which is the binding site for flagellar export chaperones in complex with their cognate F-type substrates, thereby inhibiting F-type protein export and promoting RH-type protein export. When the HBB structure is complete, FlhA_L-C_ is detached from the hydrophobic dimple and binds to the D1 and D3 domains of its nearest FlhA_C_ subunits in the FlhA_C_ ring, locking the FlhA_C_ ring structure in the F state. As a result, flagellar export chaperones in complex with their cognate F-type substrates bind to the hydrophobic dimples at a nanomolar affinity, allowing the fT3SS to efficiently transport the F-type substrates into the central channel of the growing flagellar structure and to the distal end for assembly^22,31^. Because the *flhA(G368C)* mutation in the GYXLI motif has been shown to affect chaperone binding to the conserved hydrophobic dimple of FlhA_C_^33,34^, we analyzed the export switching ability of the FlhA_C_ mutants with AAAA and GGGG mutations in the GYXLI motif. The AAAA mutation delayed substrate specificity switching of the fT3SS from the RH-type to the F-type, resulting in a polyhook-filament phenotype while the GGGG mutation totally inhibited the substrate specificity switching, causing a polyhook phenotype (Supplementary Figs. 1 and 2). The intragenic AAAV/T and GGGV suppressor mutations restored the export switching function of the fT3SS to a significant degree (Fig. 2). These results suggest that a conformational change of the GYXLI motif is required not only to release FlhA_L-C_ from the conserved hydrophobic dimple but also to remodel hydrophobic interaction networks in FlhA_C_ to create an appropriate chaperone-binding site in the dimple upon hook completion.

Residues 301–350 of the FliK_C_ core domain and the last five residues of the intrinsically disordered FliK_C_ region are important for the export switching function of FliK^39^. Photo-crosslinking experiments have shown a direct interaction between the FliK_C_ core domain and FlhB_C_^23^. The *flhB(P270A)* mutation inhibits substrate specificity switching of the fT3SS from the RH-type to the F-type at an appropriate timing of hook assembly, thereby producing polyhooks with or without filament attached^40^. The *flhB(P270A)* mutation does not affect the photo-crosslinking efficiency between FliK_C_ and FlhB_C_, indicating that FliK cannot efficiently transmit the hook length signal to the FlhA_C_ ring through an interaction between FliK and FlhB_C_ with the P270A mutation^24^. The *flhA(A489E)* suppressor mutation, located at the chaperone binding site of FlhA_C_, partially restores the motility of the *flhB(P270A)* mutant. This *flhA* mutation increases the probability of filament assembly but does not shorten the polyhook length significantly. These observations suggest that the interaction between FlhB_C_ and FlhA_C_ is critical for the initiation of F-type protein export^24^. Here, we showed that the *fliK(Q338R)* and *fliK(A405V)* mutations significantly shortened the length of polyhooks produced by the AAAA mutant, thereby improving flagella-driven motility in soft agar (Fig. 3). The *fliK(Q338R)* and *fliK(A405V)* mutations alone showed no significant motility phenotype (Supplementary Fig. 5), suggesting that these two *fliK* mutations do not inhibit or facilitate the interaction of FliK_C_ with FlhB_C_. Therefore, we propose that the *fliK(Q338R)* and *fliK(A405V)* mutations allow the FliK_C_-FlhB_C_ complex to efficiently bind to FlhA_C_ with the AAAA mutation, thereby inducing the dissociation of FlhA_L-C_ from the conserved hydrophobic dimple to terminate RH-type protein export at a more appropriate timing of hook assembly. However, these extragenic suppressor mutations in the *fliK* gene did not improve the efficiency of F-type protein export at all (Fig. 3d, 3rd and 4th rows), and these suppressor mutants produced short flagellar filaments like the AAAA mutant (Supplementary Fig. 2). Thus, it seems unlikely that the FliK_C_-FlhB_C_ complex is directly involved in the formation of the appropriate chaperone-binding site in the hydrophobic dimple of FlhA_C_ after complete cessation of RH-type protein export.

The hook length of *Salmonella* is controlled to about 55 nm with an error of about 10%. The average hook length of the ΔHI B* mutant is nearly the same as that of the wild-type strain, but the hook length distribution of this mutant is much broader than that of the wild type (Fig. 2c)^38^. Therefore, FliH and FliI are necessary for FliK to measure hook length in a more accurate manner. Here, we showed that removal of both FliH and FliI from the intragenic AAAV/T and GGGV suppressor mutants markedly reduce the substrate specificity switching efficiency of the fT3SS, resulting in much longer polyhooks (Fig. 2c). Because FlhA_L-C_ binds tightly to the hydrophobic dimple of FlhA_C_ during HBB assembly^31^, biological energy may be required for efficient dissociation of FlhA_L-C_ from the dimple. FliH, FliI, and FliJ assemble into the cytoplasmic ATPase ring complex at the base of the flagellum, and ATP hydrolysis by the FliI ATPase turns an inactive export gate complex into a highly active protein transporter through the interaction of FliJ with FlhA ^3^. Since the W354A and E351A/D356A mutations in FlhA_L-C_ inhibit the export of F-type proteins but not that of RH-type proteins, it has been proposed that the interaction between FlhA_L_ and FliJ may also be required to efficiently switch the FlhA_C_ ring structure from the RH state to the F state upon hook completion^22^. Because the AAAV/T and GGGV mutations did not inhibit the interaction of FlhA_C_ with FliJ (Fig. 5b), we propose that the efficient transition of the FlhA_C_ ring from the RH state to the F state induced by the FliJ-FlhA_L_ interaction may require energy derived from ATP hydrolysis by the cytoplasmic ATPase ring complex.

The highly conserved Tyr-106 residue of FliT is required for the interaction with FlhA_C_ (PDB ID: 6CH2)^29,30^. Comparison of the FlhA_C_ structures with and without FliT bound has shown that the binding of FliT to FlhA_C_ induces a rotation of domain D2 relative to domain D1 of FlhA_C_ through a conformational change in the GYXLI motif, thereby allowing Tyr-106 of FliT to bind efficiently to the hydrophobic dimple of FlhA_C_ (Supplementary Fig. 9). Purified FlhA_C_ with the AAAV mutation formed the nonameric ring like the wild-type FlhA_C_ (Fig. 4), suggesting that FlhA_C_ with the AAAV mutation prefers to adopt an F-type conformation in solution. However, this AAAV mutation reduced the binding affinity of FlhA_C_ for the FlgN-FlgK chaperone substrate complex (Fig. 5a, 3rd row), suggesting that this mutation restricts the rotation of domain D2 relative to domain D1. The AAAV mutation inhibited the secretion of FlgK and FliD in the absence of FliH and FliI but not in their presence (Fig. 2d and Supplementary Fig. 3). Therefore, we suggest that FlhA_C_ requires the support of FliH and FliI to maintain an appropriate conformation of the GYXLI motif to facilitate efficient docking of the chaperone-substrate complex to FlhA_C_.

In addition to flagellar export chaperones in complex with their cognate F-type substrates, RH-type substrates also bind to FlhA_C_^33,41^, and FliH and FliI are required for hierarchical targeting of export substrates and chaperone-substrate complexes to FlhA_C_^38^. Here, we found that the fT3SS with the AAAA or GGGG mutation in FlhA_C_ recognizes FlgL as an F-type substrate in the presence of FliH and FliI but as an RH-type substrate in their absence (Fig. 2d), indicating that FliH and FliI help the fT3SS with the AAAA or GGGG mutation correct substrate recognition errors that occur during flagellar assembly. Because FlhA_C_ can take different conformations through conformational changes in the GYXLI motif^34^, we propose that FliH and FliI also support FlhA in the proofreading function to bring strict order in flagellar protein export for efficient assembly of the flagellum and that a specific conformational change of the GYXLI motif is required for this proofreading function to be properly performed. RH-type substrates have a common hydrophobic sequence (FXXXΦ; Φ is a hydrophobic residue) in their N-terminal region, named gate recognition motif (GRM) responsible for an interaction with FlhB_C_. The interaction between the GRM and FlhB_C_ is essential for RH-type protein export^42–44^. Because this GRM sequence is not seen in the N-terminal region of FlgL, we conclude that the GRM is not critical for the proofreading function by FlhA_C_ with FliH and FliI.

## Methods

### Bacterial strains, plasmids, P22-mediated transduction, and media

*Salmonella* strains and plasmids used in this study are listed in Supplementary Table 1. To identify and purify extragenic suppressor mutations, P22-mediated transduction was performed using P22HT*int*^45^. L-broth contained 10 g of Bacto-Tryptone, 5 g of yeast extract and 5g of NaCl per liter. Soft tryptone agar plates contained 10 g of Bacto Tryptone, 5 g of NaCl and 0.35% (w/v) Bacto-Agar per liter. Ampicillin and tetracycline were added as needed at a final concentration of 100 μg ml^-1^ and 15 μg ml^-1^, respectively.

### DNA manipulations

DNA manipulations were performed using standard protocols. Site-directed mutagenesis was carried out using Prime STAR Max Premix as described in the manufacturer’s instructions (Takara Bio). All mutations were confirmed by DNA sequencing (Eurofins Genomics).

### Motility assays in soft agar

Fresh colonies were inoculated onto soft tryptone agar plates and incubated at 30°C. At least six measurements were carried out.

### Secretion assays

Secretion assays were carried out described previously^46^. *Salmonella* cells were grown in 5 ml of L-broth containing ampicillin with shaking until the cell density had reached an OD_600_ of ca. 1.2–1.4. Cultures were centrifuged to obtain cell pellets and culture supernatants, separately. The cell pellets were resuspended in sodium dodecyl sulfate (SDS)-loading buffer solution [62.5 mM Tris-HCl, pH 6.8, 2% (w/v) SDS, 10% (w/v) glycerol, 0.001% (w/v) bromophenol blue] containing 1 μl of 2-mercaptoethanol. Proteins in each culture supernatant were precipitated by 10% trichloroacetic acid and suspended in a Tris/SDS loading buffer (one volume of 1 M Tris, nine volumes of 1 X SDS-loading buffer solution) containing 1 μl of 2-mercaptoethanol. Both whole cellular proteins and culture supernatants were normalized to a cell density of each culture to give a constant number of *Salmonella* cells. After boiling at 95°C for 3 min, these protein samples were separated by SDS–polyacrylamide gel (normally 12.5% acrylamide) electrophoresis and transferred to nitrocellulose membranes (Cytiva) using a transblotting apparatus (Hoefer). Then, immunoblotting with polyclonal anti-FlgD, anti-FlgE, anti-FliK, anti-FlgK, anti-FlgL, anti-FlgM, anti-FliC, anti-FliD, or anti-FlhA_C_ antibody as the primary antibody and anti-rabbit IgG, HRP-linked whole Ab Donkey (GE Healthcare) as the secondary antibody was carried out. Detection was performed with an ECL prime immunoblotting detection kit (GE Healthcare). Chemiluminescence signals were detected by a Luminoimage analyzer LAS-3000 (GE Healthcare). All image data were processed with Photoshop (Adobe). At least three measurements were performed.

### Electron microscopy

HBBs and polyhook-basal bodies (PHBBs) were purified as described previously^38^. *Salmonella* cells were grown in 500 ml of L-broth containing ampicillin at 30°C with shaking until the cell density had reached an OD_600_ of ca. 1.0. After centrifugation (10,000 g, 10 min, 4°C), the cells were suspended in 20 ml of ice-cold 0.1 M Tris-HCl pH 8.0, 0.5 M sucrose, and EDTA and lysozyme were added at the final concentrations of 10 mM and 0.1 mg ml^-1^, respectively. The cell suspensions were stirred for 30 min at 4°C, and Triton X-100 and MgSO_4_ were added at final concentrations of 1% (w/v) and 10 mM, respectively. After stirring on ice for 1 hour, the cell lysates were adjusted to pH 10.5 with 5 M NaOH and then centrifuged (10,000 g, 20 min, 4°C) to remove cell debris. After ultracentrifugation (45,000 g, 60 min, 4°C), pellets were resuspended in 10 mM Tris-HCl, pH 8.0, 5 mM EDTA, 1% Triton X-100, and this solution was loaded a 20–50% (w/w) sucrose density gradient in 10 mM Tris-HCl, pH 8.0, 5 mM EDTA, 1% Triton X-100. After ultracentrifugation (49,100 g, 13 h, 4°C), HBBs and PHBBs with or without filament attached were collected and ultracentrifuged (60,000 g, 60 min, 4°C). Pellets were suspended in 50 mM glycine, pH 2.5, 0.1% Triton X100 to depolymerize the filaments. After ultracentrifugation (60,000 g, 60 min, 4°C), pellets were resuspended in 50 μl of 10 mM Tris-HCl, pH 8.0, 5 mM EDTA, 0.1% Triton X100. The HBBs and PHBBs were negatively stained with 2% (w/v) uranyl acetate. Electron micrographs were taken using JEM-1400Flash (JEOL, Tokyo, Japan) operated at 100 kV. The length of hooks and polyhooks was measured by ImageJ version 1.52 (National Institutes of Health).

### Fluorescence microscopy

*Salmonella* cells were grown in 5 ml of L-broth containing ampicillin. The cells were attached to a cover slip (Matsunami glass, Japan), and unattached cells were washed away with motility buffer (10 mM potassium phosphate pH 7.0, 0.1 mM EDTA, 10 mM L-sodium lactate). Them, flagellar filaments were labelled using anti-FliC antibody and anti-rabbit IgG conjugated with Alexa Fluor 594 (Invitrogen). After washing twice with the motility buffer, the cells were observed by an inverted fluorescence microscope (IX-83, Olympus) with a 150× oil immersion objective lens (UApo150XOTIRFM, NA 1.45, Olympus) and an Electron-Multiplying Charge-Coupled Device camera (iXon^EM^+897-BI, Andor Technology). Fluorescence images of filaments labeled with Alexa Fluor 594 were merged with bright field images of cell bodies using ImageJ software version 1.52 (National Institutes of Health).

### Purification of His-tagged wild-type FlhA_C_ and its mutant variants

His-FlhA_C_ and its mutant variants were expressed in the *E. coli* BL21 Star (DE3) strain and purified by Ni-NTA affinity chromatography with a Ni-NTA agarose column (QIAGEN), followed by size exclusion chromatography with a HiLoad 26/600 Superdex 75 pg column (GE Healthcare) at a flow rate of 2.5 ml min^-1^ equilibrated with 50 mM Tris-HCl, pH 8.0, 100 mM NaCl, 1 mM EDTA as described previously^47^.

### Analytical size exclusion chromatography

Analytical size exclusion chromatography was performed with a Superdex 75 HR 10/30 column (GE Healthcare). A 500 μl solution of purified His-FlhA_C_ and its mutant variants (10 μM) were run on the SEC column equilibrated with 50 mM Tri-HCl, pH 8.0, 150 mM NaCl at a flow rate of 0.5 ml min^-1^. Bovine serum albumin (66.4 kDa) and ovalbumin (44 kDa) were used as size markers. All fractions were run on SDS-PAGE and then analyzed by Coomassie Brilliant blue (CBB) staining.

### Far-UV CD spectroscopy

Far-UV CD spectroscopy of His-FlhA_C_ or its mutant variants was carried out at room temperature using a Jasco-720 spectropolarimeter (JASCO International Co., Tokyo, Japan) as described previously^48^. The CD spectra of His-FlhA_C_ and its mutant forms were measured in 20 mM Tris-HCl, pH 8.0 using a cylindrical fused quartz cell with a path length of 0.1 cm in a wavelength range of 200 nm to 260 nm. Spectra were obtained by averaging five successive accumulations with a wavelength step of 0.5 nm at a rate of 20 nm min^-1^, response time of 8 sec, and bandwidth of 2.0 nm.

### Pull-down assays by GST affinity chromatography

FlgK was expressed in the *E. coli* BL21 Star (DE3) cells and purified as described previously^49^. GST-FlgN and GST-FliJ were purified by GST affinity chromatography with a glutathione Sepharose 4B column (GE Healthcare) from the soluble fractions from the S*almonella* SJW1368 strain transformed with a pGEX-6p-1-based plasmid as described previously^29^. Purified His-FlhA_C_ or its mutant variants were mixed purified GST-FlgN/FlgK complex or purified GST-FliJ, and each mixture was dialyzed overnight against PBS (8 g of NaCl, 0.2 g of KCl, 3.63 g of Na_2_HPO_4_•12H_2_O, 0.24 g of KH_2_PO_4_, pH 7.4 per liter) at 4°C with three changes of PBS. A 5 ml solution of each mixture was loaded onto a glutathione Sepharose 4B column (bed volume, 1 ml) pre-equilibrated with 20 ml of PBS. After washing of the column with 10 ml PBS at a flow rate of ca. 0.5 ml min^-1^, bound proteins were eluted with 50 mM Tris-HCl, pH 8.0, 10 mM reduced glutathione. Fractions were analyzed by SDS-PAGE with CBB staining.

### High-speed atomic force microscopy

HS-AFM imaging of FlhA_C_ ring formation was carried out in solution using a laboratory-built HS-AFM^50,51^. A 2 μl solution of purified FlhA_C_ monomer (2 μM) in 50 mM Tris-HCl, pH 8.0, 100 mM NaCl was placed on a freshly cleaved mica surface attached to a cylindrical glass stage. After incubating at room temperature for 3 min, the mica surface was rinsed thoroughly with 20 μl of 10 mM Tris-HCl, pH 6.8 to remove any residual molecules. Subsequently, the sample was immersed in a liquid cell containing 60 μl of 10 mM Tris-HCl, pH 6.8. AFM imaging was performed in a tapping mode using small cantilevers (AC7, Olympus). The AFM images were recorded at 200 ms/frame in a scanning area of 100 × 100 nm^2^ with 150 × 150 pixels. For AFM image analysis, a low-pass filter was applied to remove random noise from the HS-AFM image, and a flattening filter was used to flatten the entire xy plane. Such image processing was carried out using a laboratory-developed software built on Igor Pro (HULINKS).

### Statistics and reproducibility

Statistical tests, sample size, and number of biological replicates are reported in the figure legends. Statistical analyses were done using Excel (Microsoft).

## Data availability

All data generated during this study are included in this published article, and Supplementary Information. Strains, plasmids, polyclonal antibodies, and all other data are available from the corresponding author on reasonable request.

## Acknowledgements

This work was supported in part by JSPS KAKENHI Grant Numbers JP20K15749 and JP22K06162 (to M.K.), JP19H03182, JP22H02573, and JP22K19274 (to T.M.), and JP21H01772 (to T.U.). This work has also been supported by Platform Project for Supporting Drug Discovery and Life Science Research (BINDS) from AMED under Grant Number JP19am0101117 and JP21am0101117 (to K.N.), by the Cyclic Innovation for Clinical Empowerment (CiCLE) from AMED under Grant Number JP17pc0101020 (to K.N.), and by JEOL YOKOGUSHI Research Alliance Laboratories of Osaka University (to K.N.).

## Author Contributions

T.M. and K.N. conceived and designed research; M.K., T.M., and T.U. preformed research; M.K., T.M., and T.U. analysed the data; and T.M. and K.N. wrote the paper based on discussion with other authors.

