## Supplementary information for "FliH and FliI help FlhA bring strict order to flagellar protein export in *Salmonella*"

**FlhA, together with the flagellar ATPase complex, brings strict order to flagellar protein export in *Salmonella***

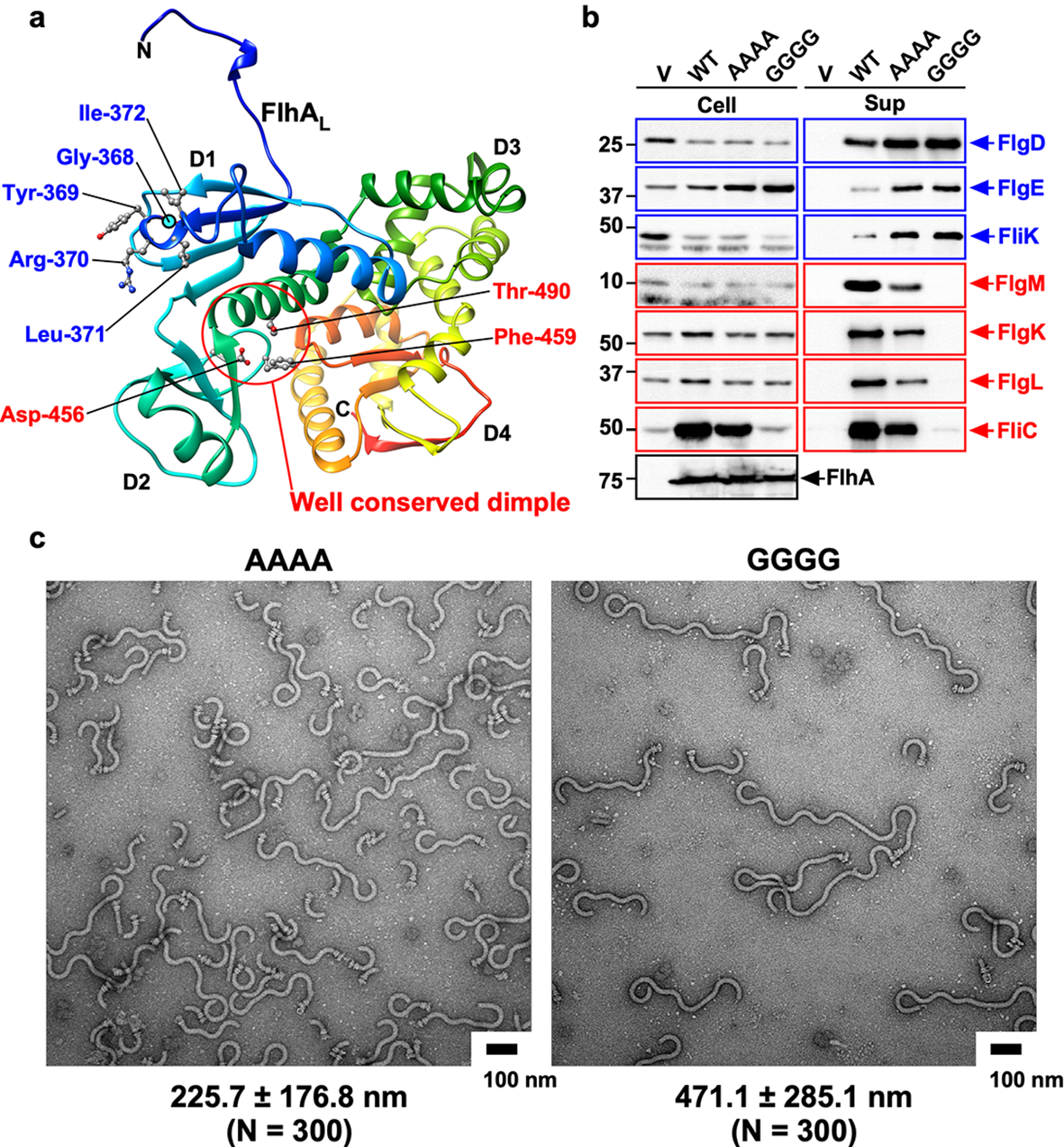

**Supplementary Fig. 1. Effect of the AAAA and GGGG mutations on flagellar protein export and assembly. (a)** Structural model of the FlhA_C_ monomer (PDB ID: 3A5I). FlhA_C_ consists of four domains, D1, D2, D3 and D4, and a flexible linker (FlhA_L_). A highly conserved Gly-368 residue (cyan circle) in domain D1 forms the conserved GYXLI motif along with Tyr-369, Arg-370, Leu-371, and Ile-372, which forms a short α-helix. The well-conserved dimple including the Asp-456, Phe-459, and Thr-490 residues is responsible for the interaction of FlhA_C_ with flagellar export chaperones in complex with F-type substrates. The Cα backbone is color-coded from blue to red, going through the rainbow colors from the N- to the C-terminus. **(b)** Secretion analysis of FlgD, FlgE, FliK, FlgM, FlgK, FlgL, and FliC by immunoblotting. Whole cell proteins (Cell) and culture supernatants (Sup) were prepared from the *Salmonella* NH001 (∆*flhA*) strain transformed with pTrc99AFF4 (indicated as V), pMM130 (indicated as WT), pMKM130-A4 (indicated as AAAA), or pMKM130-G4 (indicated as GGGG). A 3 μl solution of each sample normalized to an optical density of OD_600_ was subjected to SDS-PAGE and analyzed by immunoblotting using polyclonal anti-FlgD (1st row), anti-FlgE (2nd row), anti-FliK (3rd row), anti-FlgM (4th row), anti-FlgK (5th row), anti-FlgL (6th row), anti-FliC (7th row), or anti-FlhA_C_ (8th row) antibody. RH-type and F-type substrates are highlighted in blue and red, respectively. Molecular mass markers (kDa) are shown on the left. **(c)** Electron micrographs of polyhook-basal bodies isolated from the AAAA and GGGG mutants. The average polyhook length and standard deviations are shown. N indices the number of polyhook-basal bodies that were measured.

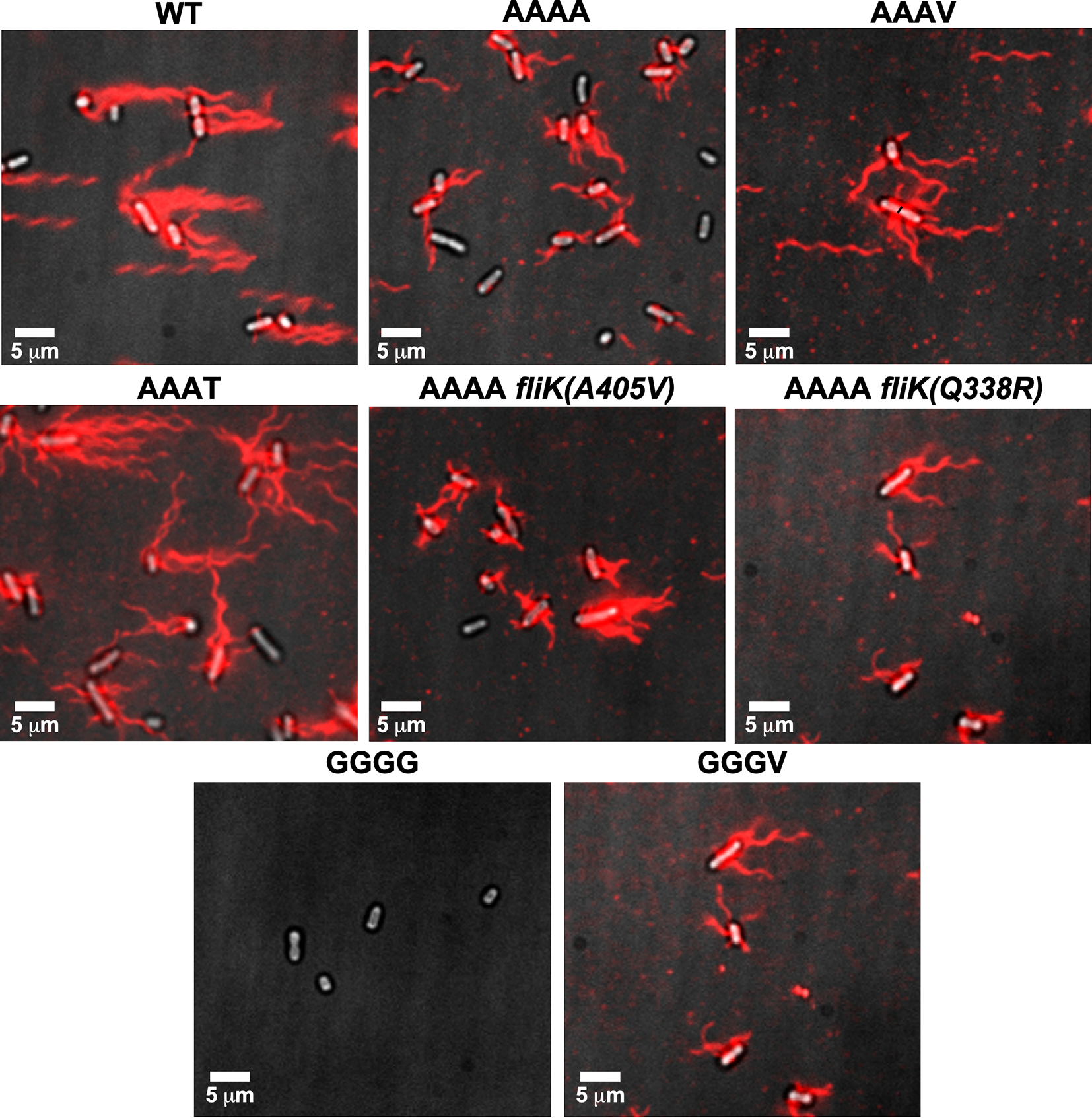

**Supplementary Fig. 2**. **Effect of suppressor mutations on flagellar filament formation in the AAAA and GGGG mutants.** Fluorescent images of NH001 carrying pMM130 (indicated as WT), MMA130A4 (indicated as AAAA), MMA130A4-5 (indicated as AAAV), MMA130A4-7 (indicated as AAAT), MMA130A4-3 [AAAA *fliK(A405V)*], MMA130A4-10 [AAAA *fliK(Q338R)*], MMA130G4 (indicated as GGGG), or MMA130G4-3 (indicated as GGGV). Fresh colonies were grown in L-broth containing ampicillin until the cells reached the stationary phase, and then flagellar filaments were labelled with a fluorescent dye, Alexa Fluor 594. The fluorescence images of the filaments labelled with Alexa Fluor 594 (red) were merged with the bright field images of the cell bodies.

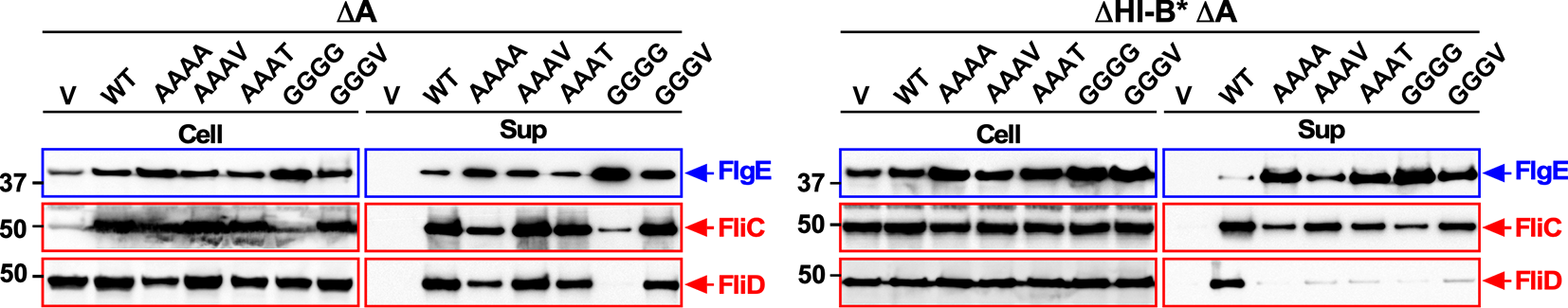

**Supplementary Fig. 3. Protein transport properties of the AAAV/T and GGGV mutants isolated as intragenic suppressor mutants from the AAAA and GGGG mutants, respectively, in the presence (left panels) and absence (right panels) of FliH and FliI.** Immunoblotting using polyclonal anti-FlgE (1st row), anti-FliC (2nd row), or anti-FliD (3rd row) antibody, of whole cell proteins (Cell) and culture supernatants (Sup) prepared from the *Salmonella* NH001 (∆*flhA*, indicated as ∆A) or NH003 [∆*fliH-fliI flhB(P28T)* ∆*flhA*, indicated as ∆HI-B* ∆A] strain transformed with pTrc99AFF4 (indicated as V), pMM130 (indicated as WT), pMKM130-A4 (indicated as AAAA), pMKM130-A3V (indicated as AAAV), pMKM130-A3T (indicated as AAAT), pMKM130-G4 (indicated as GGGG), or pMKM130-G3V (indicated as GGGV). RH-type and F-type substrates are highlighted in blue and red, respectively. Molecular mass markers (kDa) are shown on the left.

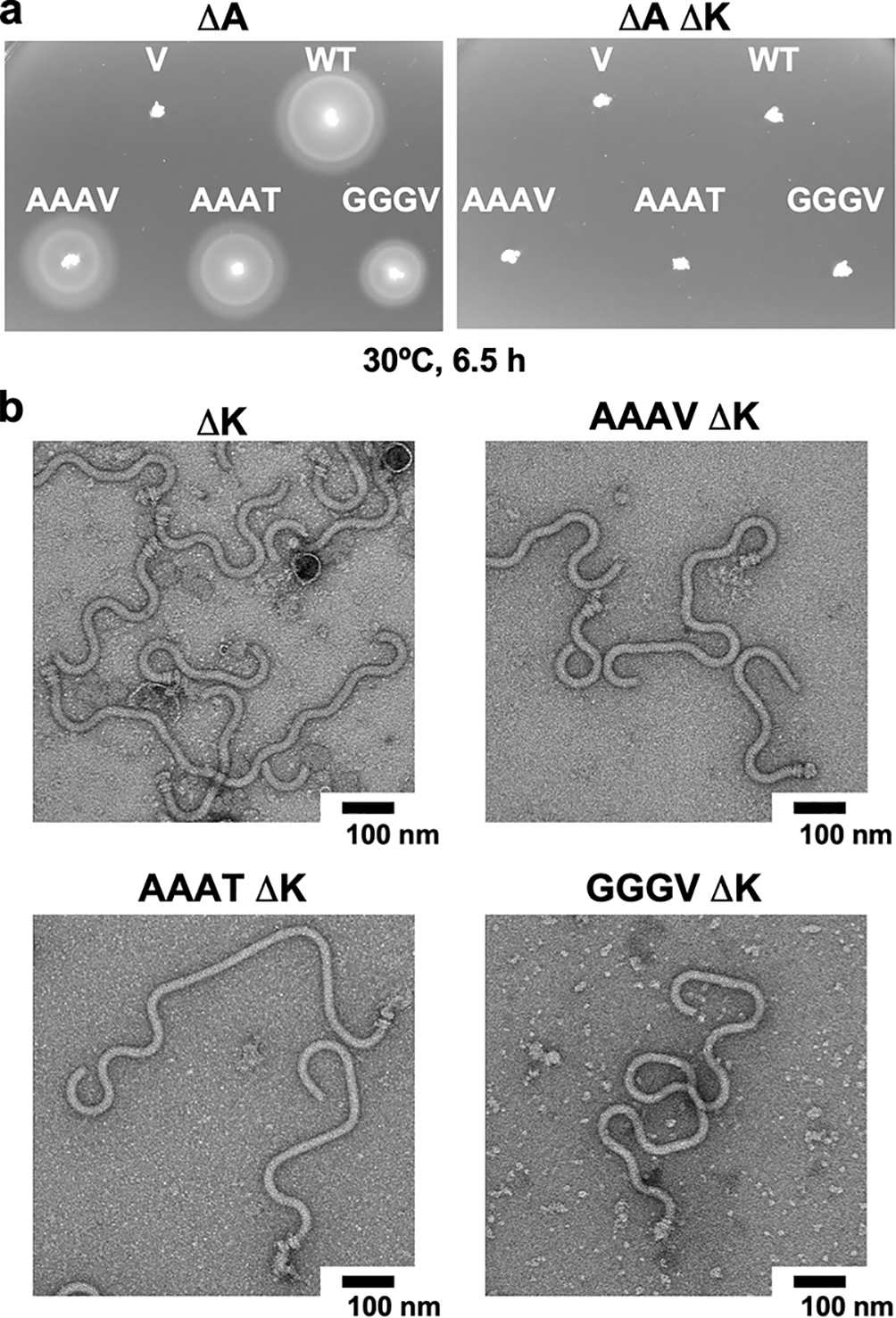

**Supplementary Fig. 4. Effect of FliK deletion on the export switching function of the AAAV/T and GGGV mutants.** **(a)** Motility of the *Salmonella* NH001 (*∆flhA*, indicated as *∆*A) (left panel) and NH001iK (∆*flhA* ∆*fliK*::*tetRA*, indicated as *∆*A *∆*K) (right panel) strains carrying pTrc99AFF4 (V), pMM130 (WT), pMKM130-A3V (AAAV), pMKM130-A3T (AAAT), or pMKM130-G3V (GGGV) in soft agar. Plates were incubated at 30ºC for 6.5 hours. **(b)** Electron micrographs of polyhook-basal bodies isolated from the above transformants.

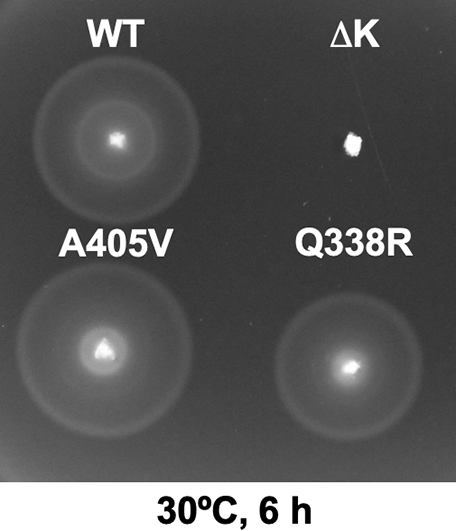

**Supplementary Fig. 5. Effect of second site *fliK* mutations on motility.** Motility of SJW1103 (wild-type, indicated as WT), TH8426 (∆*fliK*, indicated as ∆K), MMK130-3 [*flhK(A405V)*, indicated as A405V], or MMK130-10 [*fliK(Q338R)*, indicated as Q338R] in soft agar. Plates were incubated at 30ºC for 6 hours.

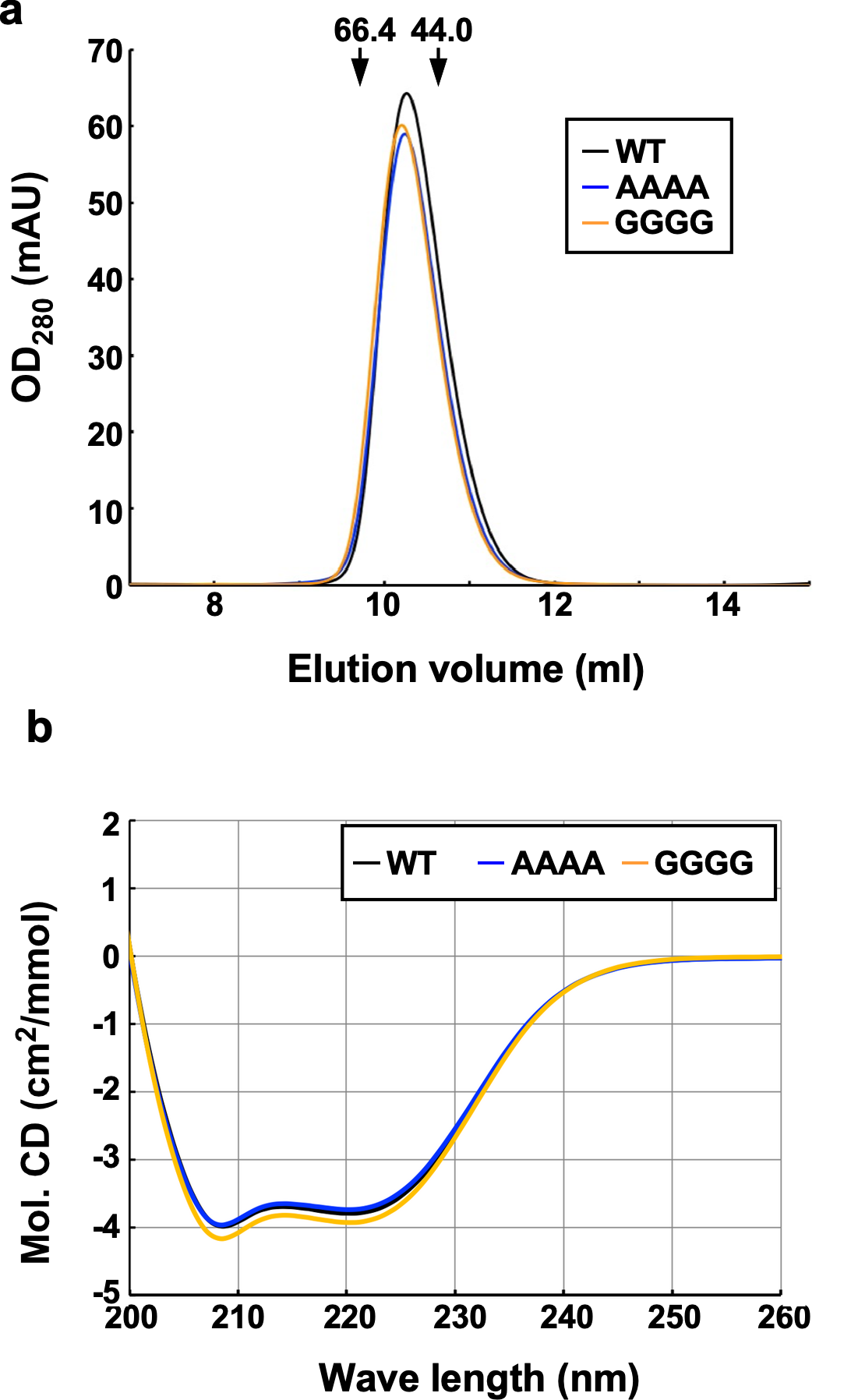

**Supplementary Fig. 6**. Effect of the AAAA and GGGG mutations on the monomeric FlhA_C_ conformation. **(a)** Effect of the AAAA and GGGG mutations on hydrodynamic properties of FlhA_C_. A 500 μl solution of each purified sample (10 μM) was run on a Superdex 75HR 10/30 column equilibrated with 50 mM Tri-HCl, pH 8.0, 150 mM NaCl. The elution peaks of His-FlhA_C_ (WT, black), His-FlhA_C-AAAA_ (AAAA, blue), and His-FlhA_C-GGGG_ (GGGG, orange) are 10.3 ml, 10.2 ml, and 10.2 ml, respectively. Arrows indicate the elution peaks of bovine serum albumin (66.4 kDa) and ovalbumin (44 kDa), which are 9.7 ml and 10.7 ml, respectively. **(b)** Effect of the AAAA and GGGG mutations on far-UV CD spectra of FlhA_C_. Measurements were carried out at room temperature in 20 mM Tris-HCl, pH 8.0, in a quartz cell with a path length of 1 mm.

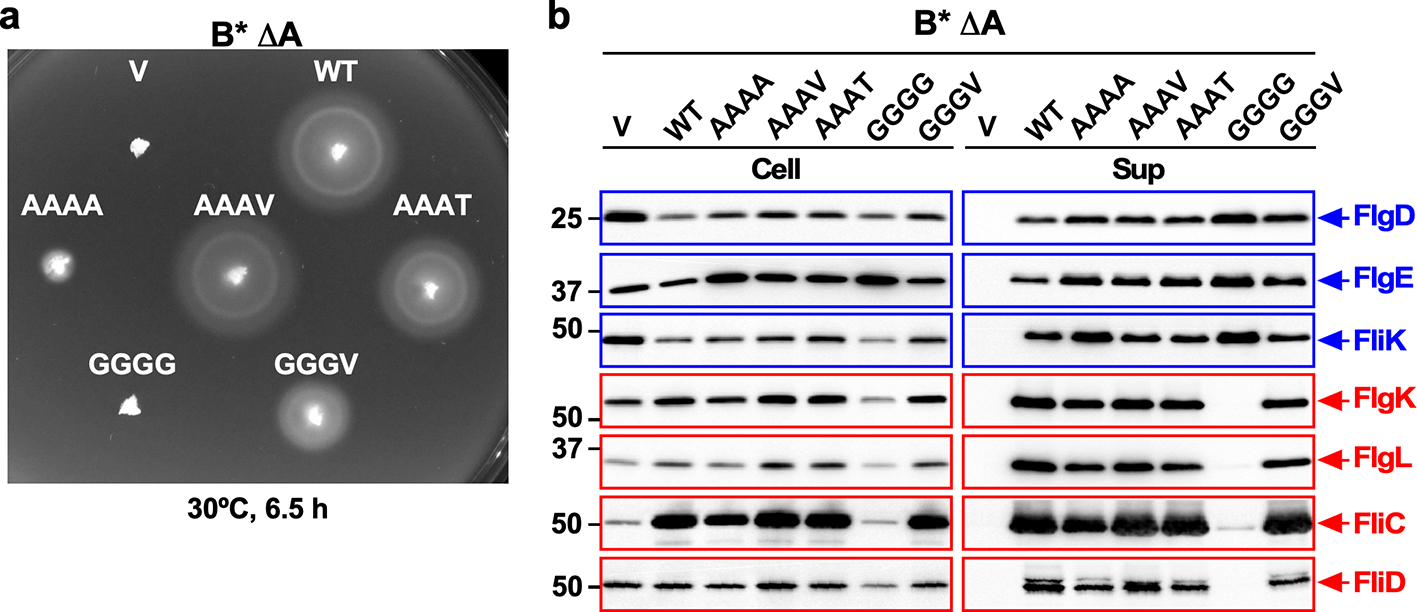

**Supplementary Fig. 7**. **Effect of the *flhB(P28T)* mutation on flagellar protein export by the AAAV/T and GGGV mutants. (a)** Motility of the *Salmonella* NH002 [*flhB(P28T)* *∆flhA*, indicated as B* *∆*A] strain carrying pTrc99AFF4 (V), pMM130 (WT), pMKM130-A4 (AAAA), pMKM130-A3V (AAAV), pMKM130-A3T (AAAT), pMKM130-G4 (GGGG), or pMKM130-G3V (GGGV) in soft agar. Plates were incubated at 30ºC for 6.5 hours. **(b)** Immunoblotting using polyclonal anti-FlgD (1st row), anti-FlgE (2nd row), anti-FliK (3rd row), anti-FlgK (4th row), anti-FlgL (5th row), anti-FliC (6th row), or anti-FliD (7th row) antibody, of whole cell proteins (Cell) and culture supernatants (Sup) prepared from the above transformants. RH-type and F-type substrates are highlighted in blue and red, respectively. The positions of molecular mass markers are indicated on the left.

**Supplementary Fig. 8**. **Effect of FlgN deletion on flagellar protein export by the AAAA and GGGG mutants in the absence of FliH and FliI.** Immunoblotting, using polyclonal anti-FlgD (1st row) and anti-FlgL (2nd row), of whole cell proteins and culture supernatant fractions prepared from the *Salmonella* NH003gN [∆*fliH-fliI flhB(P28T)* *∆flhA* ∆*flgN*::*tetRA*, indicated as *∆*HI-B* *∆*A *∆*N] strain transformed with pTrc99AFF4 (V), pMM130 (WT), pMKM130-A4 (AAAA), or pMKM130-G4 (GGGG). The positions of molecular mass markers are indicated on the left.

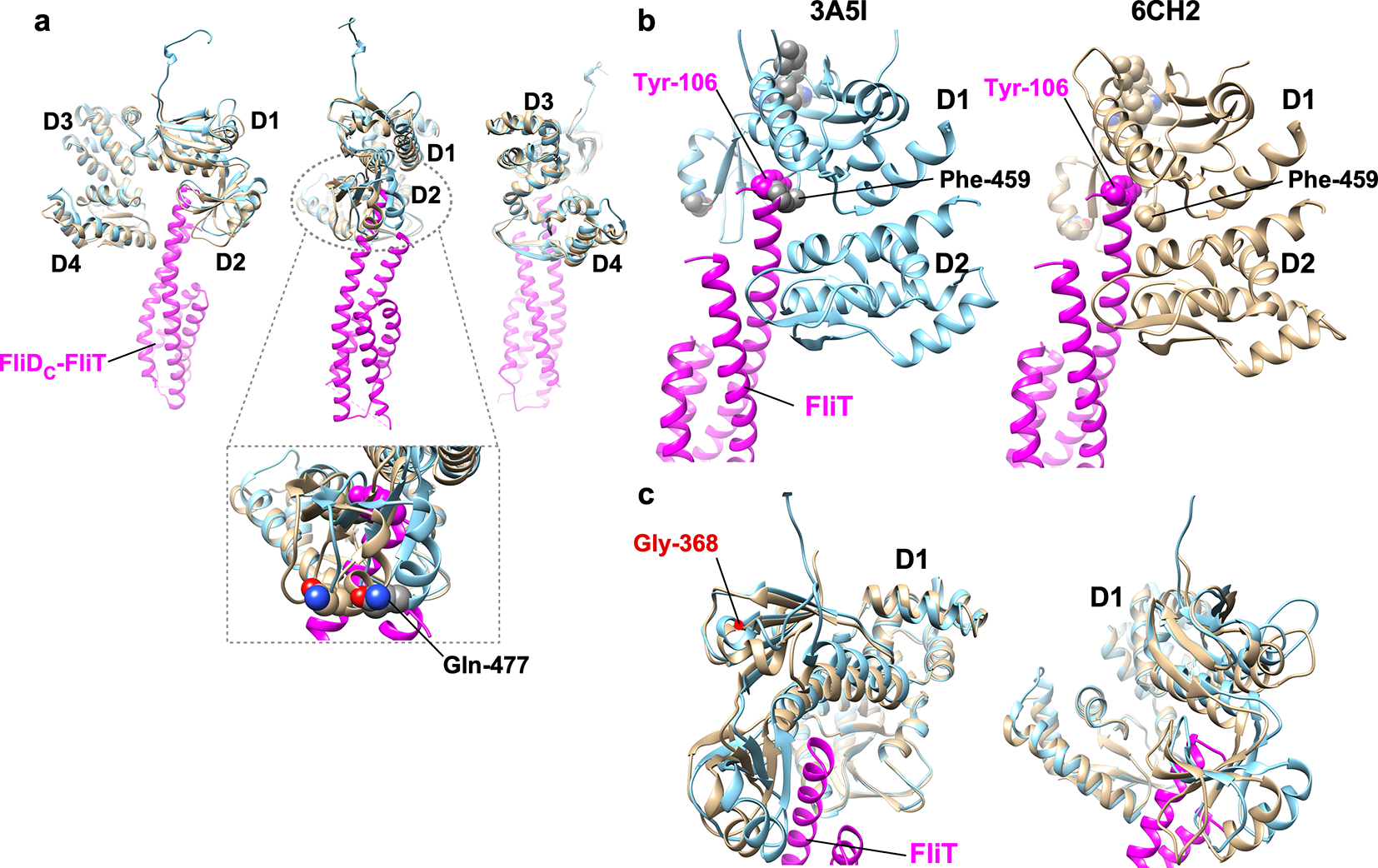

**Supplementary Fig. 9**. **Structural comparison of FlhA_C_ with and without the FliT chaperone.** Ribbon representations of the crystal structures of FlhA_C_ in an open conformation (sky blue) (PDB ID: 3A5I) and FlhA_C_ (tan) in complex with FliT fused with the C-terminal region of FliD (magenta) (PDB ID: 6CH2). (a) Three different views showing the FliT chaperone binds to the conserved hydrophobic dimple between domains D1 and D2 of FlhA_C_, resulting in a rotation of domain D2 relative to domain D1 together with a conformational change of the GYXLI motif coupled with a downward shifts of two α-helices of domain D1. Gln-477 is a good indicator of the rotation of domain D2 relative to domain D1. The middle and right panels are rotated 90 degrees clockwise and counterclockwise from the left panel, respectively. (b) FliT binds to a well-conserved hydrophobic dimple of FlhA_C_ located at the interface between domains D1 and D2, and the highly conserved Tyr-106 residue of FliT and the highly conserved Phe-459 residue of FlhA_C_ are directly involved in the interaction between FlhA_C_ and FliT. The rotation of domain D2 relative to domain D1 allows Tyr-106 of FliT to fit nicely into the hydrophobic dimple of FlhA_C_. (c) Gly-368 of FlhA_C_ is located within the conserved GYXLI motif of FlhA_C_. The YXLI sequence forms a short α-helix. The binding of FliT to FlhA_C_ induces a conformational change of the conserved GYXLI motif of FlhA_C_. The right panel is rotated 60 degrees clockwise from the left panel.

**Supplementary Table 1. Strains and plasmids used in this study.**

| **Strain/Plasmid** | **Relevant characteristics** | **References** |
| --- | --- | --- |
| ***E. coli*** |  |  |
| BL21 Star (DE3) | Overexpression of proteins | Novagen |
| ***Salmonella*** |  |  |
| SJW1103 | Wild-type for motility and chemotaxis | 52 |
| SJW1368 | ∆*cheW–flhD* | 53 |
| NH001 | ∆*flhA* | 36 |
| NH002 | *flhB(P28T)* *∆flhA* | 36 |
| NH003 | ∆*fliH-fliI flhB(P28T)* *∆flhA* | 36 |
| TH8426 | ∆*fliK* | 39 |
| NH001iK | ∆*flhA* ∆*fliK*::*tetRA* | This study |
| NH003gN | ∆*fliH-fliI flhB(P28T)* *∆flhA* ∆*flgN*::*tetRA* | This study |
| MMA130A4 | NH001 harboring pMKM130-A4 | This study |
| MMA130A4-3 | NH001 harboring pMKM130-A4 *fliK(A405V)* | This study |
| MMA130A4-5 | NH001 harboring pMKM130-A3V | This study |
| MMA130A4-7 | NH001 harboring pMKM130-A3T | This study |
| MMA130A4-10 | NH001 harboring pMKM130-A4 *fliK(Q338R)* | This study |
| MMA130G4 | NH001 harboring pMKM130-G4 | This study |
| MMA130G4-3 | NH001 harboring pMKM130-G3V | This study |
| MMK130-3 | *fliK(A405V)* | This study |
| MMK130-10 | *fliK(Q338R)* | This study |
| **Plasmids** |  |  |
| pTrc99AFF4 | Modified pTrc expression vector | 54 |
| pGEX-6p-1 | Expression vector | GE Healthcare |
| pMKGK2 | pTrc99A/ FlgK | 48 |
| pMM104 | pET19b/ His-FlhA_C_ (residues 211–692) | 41 |
| pMM130 | pTrc99AFF4/ FlhA | 55 |
| pMKM130-A4 | pTrc99AFF4/ FlhA(T369A/R370A/L371A/I372A) | 34 |
| pMKM130-G4 | pTrc99AFF4/ FlhA(T369G/R370G/L371G/I372G) | 34 |
| pMMGN101 | pGEX-6p-1/ GST-FlgN | 28 |
| pMMJ1001 | pGEX-6p-1/ GST-FliJ | 35 |
| pMKM104-A4 | pET19b/ His-FlhA_C_(T369A/R370A/L371A/I372A) | This study |
| pMKM104-A3V | pET19b/ His-FlhA_C_(T369A/R370A/L371A/I372V) | This study |
| pMKM104-G4 | pET19b/ His-FlhA_C_(T369G/R370G/L371G/I372G) | This study |
| pMKM104-G3V | pET19b/ His-FlhA_C_(T369G/R370G/L371G/I372V) | This study |
| pMKM130-A3V | pTrc99AFF4/ FlhA(T369A/R370A/L371A/I372V) | This study |
| pMKM130-A3T | pTrc99AFF4/ FlhA(T369A/R370A/L371A/I372T) | This study |
| pMKM130-G3V | pTrc99AFF4/ FlhA(T369G/R370G/L371G/I372V) | This study |
